# Multi-Response Phylogenetic Mixed Models: Concepts and Application

**DOI:** 10.1101/2022.12.13.520338

**Authors:** Ben Halliwell, Barbara R. Holland, Luke A. Yates

## Abstract

The scale and resolution of trait databases and molecular phylogenies is increasing rapidly. These resources permit many open questions in comparative biology to be addressed with the right statistical tools. Multi-response (MR) phylogenetic mixed models (PMM) offer great potential for multivariate analyses of trait evolution. While flexible and powerful, these methods are not often employed by researchers in ecology and evolution, reflecting a specialised and technical literature that creates barriers to usage for many biologists. Here we present a practical and accessible guide to MR-PMM. We begin with a review of single-response (SR) PMM to introduce key concepts and outline the limitations of this approach for characterizing patterns of trait co-evolution. We emphasise MR-PMM as a preferable approach to analyses involving multiple species traits, due to the explicit decomposition of trait covariance across levels in the model hierarchy. We discuss multilevel distributional models, meta-analyses, multivariate models of evolution, and extensions to non-Gaussian response traits. We highlight techniques for causal inference using precision matrices, as well as advanced topics including prior specification and latent factor models. Using simulated data and visual examples, we discuss interpretation, prediction, and model validation. We implement many of the techniques discussed in example analyses of plant functional traits to demonstrate the general utility of MR-PMM in handling complex real world datasets. Finally, we discuss the emerging synthesis of comparative techniques made possible by MR-PMM, highlight strengths and weaknesses, and offer practical recommendations to analysts. To complement this material, we provide extensive online tutorials including side-by-side model implementations in two popular R packages, `MCMCglmm` and `brms`.

## 1 Introduction

The motivation for developing multivariate phylogenetic comparative methods is now broadly appreciated (Adams and Collyer 2018; Uyeda et al. 2015, 2018; Garamszegi 2014). Modern approaches aim to move beyond phylogenetic regression and variance partitioning of individual Gaussian variables (e.g., Lynch 1991; Pagel 1999; Blomberg et al. 2003), to methods capable of evaluating the strength, direction, and conservatism of eco-evolutionary relationships within networks of continuous and discrete variables (Westoby et al., 2023; Haba and Kutsukake, 2019; Brommer et al., 2019; Hadfield, 2010). These multivariate techniques are applicable to a broad range of species traits, from morphology, physiology, and behavior, to environmental tolerance limits and niche characteristics. Furthermore, as the tide of 21st-century data collection erodes historical constraints on the scale and complexity of analyses, opportunities to apply these methods are increasing: Advances in climate modeling, remote sensing technology, and collaborative data projects are accelerating the availability and resolution of suitable datasets (Green et al., 2022; Herberstein et al., 2022; Falster et al., 2021; Kattge et al., 2020); the genomic revolution continues to provide more accurate and complete phylogenies (Young and Gillung, 2020; Laumer et al., 2019; Smith and Brown, 2018; Yeates et al., 2016); and the computational resources necessary to pursue fully Bayesian analyses are now accessible to most researchers. Despite this coalescence of opportunities, challenges in implementing modern methods create barriers to usage for many biologists.

A robust literature developing statistical models of multivariate trait evolution has existed for some time. However, new techniques are often slow to absorb into research practice, especially where they demand a technical or conceptual leap from users. Development of the generalised least-squares framework, helped to bring phylogenetic regression and correlative analyses of trait evolution into the mainstream (Rohlf, 2001; Martins and Hansen, 1997; Lynch, 1991; Grafen, 1989; Felsenstein, 1985). Subsequent extensions to phylogenetic generalised linear mixed models (PMM) provided additional benefits, including support for non-Gaussian traits, flexibility in the specification of hierarchical group effects, and a growing familiarity with mixed models among researchers in ecology and evolution (Ives and Helmus, 2011; Ives and Garland Jr, 2010; Bolker et al., 2009; Housworth et al., 2004). Currently, PMM also supports multi-response (MR) analyses, facilitating the estimation (and decomposition) of covariances between discrete and continuous species traits (Hadfield, 2010; Hadfield and Nakagawa, 2010).

The development of user-friendly software for fitting MR-PMM (Bürkner, 2017; Hadfield, 2010) has prompted researchers from diverse scientific fields to implement the method for sophisticated comparative analyses. For example, MR-PMM has been used in anthropology to examine evolutionary relationships between climate and cranial form among neo-lithic humans (Katz et al., 2016), and the influence of phylogenetic history on marriage patterns among modern human populations (Minocher et al., 2019); in animal behaviour to study the evolution of multivariate behavioral repertoires in flies (Hernández et al., 2021), and the evolution of group size and social complexity in birds (Downing et al., 2020); in epidemiology and disease ecology to understand relationships between growth rate, transmission mode, and virulence in pathogens (Leggett et al., 2017), the role of sexual dimorphism in susceptibility to viral infection (Roberts and Longdon, 2023), and covariance in macro- and micro-parasite species richness among host species (Gutiérrez et al., 2019); and, of course, in evolutionary ecology and eco-physiology to understand the multivariate evolution of species functional traits. Examples of the latter include coevolution among milk macro-nutrient concentrations in mammals (Blomquist, 2019), interactions between climate and functional traits on plant vital rates and life history evolution (Kelly et al., 2021), the evolutionary coordination of plant hydraulic traits (Sanchez-Martinez et al., 2020), and covariance between species traits and flammability in fire-prone grasslands (Simpson et al., 2016).

Many early applications of MR-PMM to comparative biology involved critical tests of theory made possible by the estimation of phylogenetic covariances (Section 4.1). For example, the effects of sexual selection on brain and body size evolution (García-Peña et al., 2013), whether character displacement is driven by trait divergence in allopatry or sympatry (Tobias et al., 2014), and whether low rates of extra-pair paternity, long lifespans, and tolerance of harsh environments represent causes or consequences of transitions to cooperative breeding (Cornwallis et al., 2010; Downing et al., 2015; Cornwallis et al., 2017). More recently, MR-PMM has been highlighted as a powerful framework for eco-evolutionary studies incorporating spatio-temporal random effects (Gomes et al., 2023), trait-based models of community assembly (Gallinat and Pearse, 2021), and analyses of function-valued traits (e.g., parameters of species reaction norms) (Pottier et al., 2024), as well as for disentangling correlations between response variables that manifest at different levels within hierarchical datasets (Westoby et al. 2023, also see Downs and Dochtermann 2014). These strengths of MR-PMM echo a growing consensus that multivariate statistical approaches are often valuable even when response correlations are not of specific interest; fitting multiple correlated response variables allows for the inclusion of partially missing data and may improve predictive accuracy due to the sharing of information across common grouping variables (see Section 5, also see Riley et al. 2017; Pottier et al. 2024).

Despite these benefits, and the example of pioneering researchers, MR-PMM remains underutilized, even for datasets to which it would be well suited, such as recent large-scale comparative analyses of species traits (e.g., Cássia-Silva et al. 2020; Grossnickle 2020; Bruelheide et al. 2018; Díaz et al. 2016). A sobering list of publications highlighting widespread misconceptions about phylogenetic comparative methods, erroneous statistical practices, and the potential for spurious inference, confirm the need for more translational research (Uyeda et al., 2018; Cooper et al., 2016; Uyeda et al., 2015; Revell, 2010; Freckleton, 2009; Revell et al., 2008). In particular, practical guidance for biologists wanting to apply multivariate methods that provide a more meaningful decomposition of trait relationships is needed to advance the field beyond the limits of univariate approaches (Westoby et al., 2023).

This review aims to bridge the gap between theory and practice in mixed model analyses of trait evolution. We focus on Bayesian implementations of MR-PMM, because this model class provides a highly flexible framework with general utility for comparative biology, although maximum likelihood implementations do exist for certain special cases (Mitov et al., 2020; Butler et al., 2023). The capacity for MR-PMM to provide a more informative analysis of trait (co)evolution compared with SR-PMM, is well appreciated (Housworth et al., 2004; Hadfield and Nakagawa, 2010). However, despite a long history of use among quantitative geneticists (Meyer, 1991), we believe a lack of familiarity, doubts about data requirements, and practical barriers to implementation, continue to prevent common usage of MR-PMM among ecologists and evolutionary biologists. Thus, we begin section 2 with a brief conceptual introduction to mixed models. We define the basic SR-PMM, the different parametrizations used to quantify phylogenetic signal, and the implementation of alternative models of evolution (i.e., other than pure Brownian Motion). After outlining the limitations of SR models for characterising patterns of coevolution between traits, we shift focus to MR-PMM. In particular, we highlight the different components of trait covariance that are modeled in MR-PMM (e.g., phylogenetic and residual), how these covariance structures are specified using covariance matrices, and how these matrices are parameterised to estimate correlations between species traits at different levels in the model hierarchy. We discuss extensions to the basic MR-PMM including multilevel distributional models, meta-analyses, multivariate models of trait evolution, non-Gaussian response traits, and techniques for causal inference using precision matrices and graphical models. Finally, we explore a number of extended topics to showcase emerging techniques, including prior specification for shrinkage of target parameters, as well as latent factor methods for dimension reduction. In section 4 we discuss interpretation and explore data simulated from MR-PMM to clarify the distinction between phylogenetic and non-phylogenetic components of trait correlation. In Section 5 we cover predictive assessment, including methods based on posterior predictive distributions and leave-one-out (LOO) cross-validation (CV). In section 6, we present an example analysis on leaf functional traits in Eucalyptus from the AusTraits database (Falster et al., 2021). Finally, in section 8 we discuss the strengths and weaknesses of MR-PMM, highlight emerging syntheses facilitated by the method, and provide practical advice for analysts.

To support our exposition, we provide extensive GitHub tutorials, including annotated code and mathematical appendices. These tutorials demonstrate how to simulate multivariate trait data containing phylogenetic structure, and fit corresponding MR-PMM in two popular R packages, `MCMCglmm` (Hadfield, 2010) and `brms` (Bürkner, 2017). They also cover many common tasks and challenges in the MR-PMM workflow, including data cleaning and manipulation, via example analyses of publicly available, real world datasets.

## 2 Models

### 2.1 Mixed Models

Biologists are familiar with structured sampling designs: these apply when each observation from a survey or experiment is a member of one or more recognized groups. Acknowledging structure in our data is important because observations within a group will often be more similar than can be explained by available predictors. In a simple case, the hierarchical structure may be limited to a single random (or group-level) intercept. For the popular lme4 package in R, the syntax used to express this model would be:

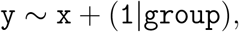

where y is the observed response, the measured covariate x specifies a fixed effect, and (1|group) specifies a random intercept (a random effect that adjusts the baseline mean) at the group level. In other words, we model y as a linear combination of effects, including a random effect that accounts for the fact that observations belong to distinct groups. A more general mathematical form for mixed models is given by,

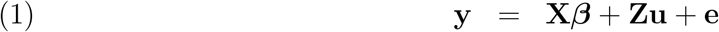

where **y** is a vector of observations. The design matrices **X** and **Z** relate fixed and random predictors to the data, the corresponding parameter vectors ***β*** and **u** contain the fixed and random effects to be estimated, and **e** is a vector of residual errors.

For the coded model formula, the random intercept (1|group) assumes the distribution of the random effect **u** is characterised by an identity matrix **I** (a matrix with 1 ‘s along the diagonal and 0’s in all off-diagonals) equal in dimension to the number of groups (where 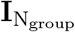 is of dimension N_group_ *×* N_group_), and scaled by a group level variance, 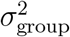. For example, for a study including data on 5 different species,

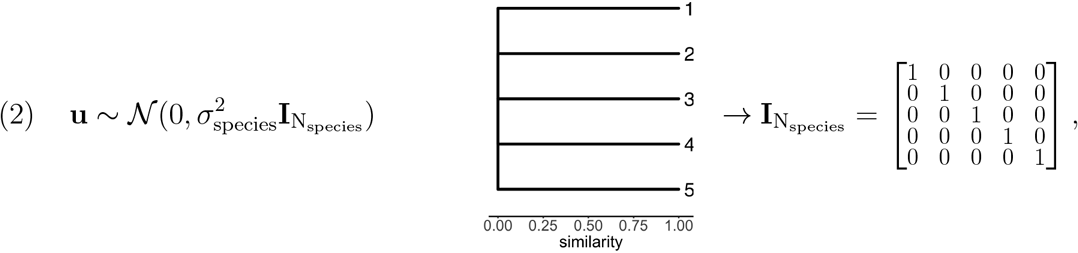

where the notation ∼ means “is distributed as” and *N* (***μ*, Σ**) denotes a (multivariate) normal distribution with mean ***μ*** and (co)variance **Σ**. The off-diagonal elements of 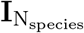 are all zero which implies independence between different levels of the grouping factor, such that no pair of species is expected to produce more similar observations than any other. Graphically, an identity matrix can be represented as a comb where tips (group levels) have no shared edges (2). The residual errors **e** are also assumed to be independent *a priori*, capturing observation-specific deviations from the modelled mean,

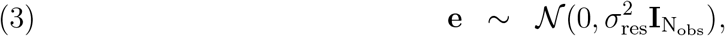

Mixed models permit the specification of complex variance structures using multiple random effects which may be given as nested (e.g., populations within species), crossed to act independently, or encoded with additional structure such as spatial or temporal relatedness. For example, if species are not uniformly distributed across the landscape, we may consider modeling spatial autocorrelation in species effects. This is achieved by adjusting our assumptions about the distribution of random effects **u**. For example, by substituting the identity matrix with a covariance matrix derived from the pairwise distances between sampling sites. Structured random effects are employed in this way to model dependencies arising from a range of data-generating processes, including evolutionary processes of species diversification.

### 2.2 Phylogenetic Mixed Models

In analyses of inter-species data, dependence arises from phylogenetic signal: the tendency for closely related species to resemble each other due to the hierarchical evolutionary history of life (Felsenstein, 1985; Pagel, 1999). This phylogenetic structure is often represented by a phylogenetic correlation matrix **C** = (*c*_*ij*_)—an N_species_ *×* N_species_ matrix that encodes our expectation of similarity among species phenotypes, where for a pair of species *i* and *j*, the expected correlation is given by the off-diagonal element *c*_*ij*_. In PMM, **C** enters the model as the expected correlation of a species-level random effect scaled by the phylogenetic variance 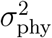. For example, given a phylogeny of 5 species, we may write

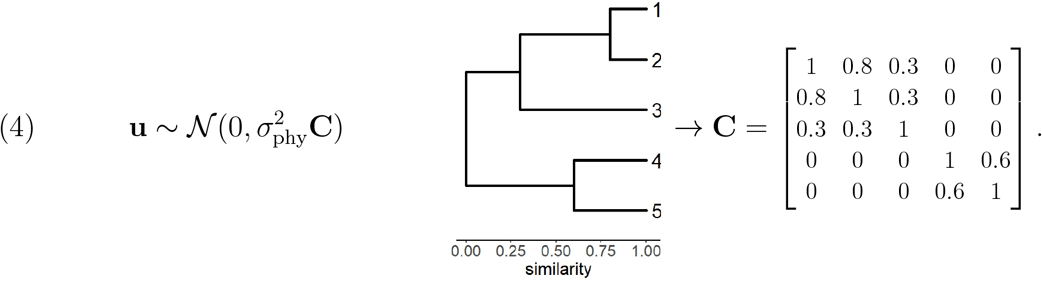

Importantly, **C** is derived by assuming a particular model of evolutionary trait change along the branches of a phylogeny (Eq. 4 assumes Brownian Motion (BM), but see Section 2.2.2). **C** is typically supplied by the user, although joint inference of the phylogenetic tree and the phylogenetic mixed model is possible in some instances, e.g., using `BEAST` (Hassler et al. 2023; Bouckaert et al. 2019). In a Bayesian setting, phylogenetic uncertainty can also be incorporated by combining posterior samples across a set of candidate topologies (Villemereuil et al. 2012; also see Nakagawa and de Villemereuil 2019).

To summarise, PMM is a special case of a linear mixed model (1) that incorporates phylogenetic information via an associated correlation matrix, **C**. Species trait values **y** are modelled as the sum of fixed effects, a phylogenetic random effect, and residual error (see Section 3.0.5 for non-Gaussian extensions). Additional random effects are permitted to model other sources of dependence among observations, such as study effects and within species replication (see Section 3.0.1).

#### 2.2.1 Phylogenetic signal

A common objective of phylogenetic comparative analyses is to quantify phylogenetic signal in species traits. Numerous metrics have been developed for this purpose, each with different assumptions and applications (Münkemüller et al., 2012). In PMM, phylogenetic signal (*λ*) describes the proportion of variance in **y** attributable to the phylogenetic structure and is expressed in terms of the estimated variance components as

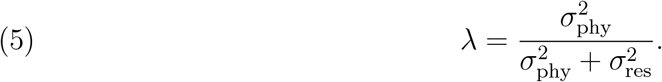

Additional variance components may be included in the denominator of (5) when random effects capture meaningful components of within- and between-species phenotypic variance, e.g., trait (co)variance owing to (shared) environmental effects (Kruuk and Hadfield, 2007). When fixed or random variance components are excluded from the denominator, (5) expresses phylogenetic signal conditional on those factors. This is generally desirable for covariates, which are often introduced to account for known biases or aspects of experimental design and thus do not contribute to the variation we seek to decompose (de Villemereuil et al., 2018). For example, when analysing data on morphometric measurements of preserved museum specimens, we may include storage duration as a fixed effect covariate to account for differences in tissue shrinkage among specimens in the collection. Such sources of variation may be considered extraneous to the estimation of *λ*, and thus excluded from the denominator of (5).

#### 2.2.2 Models of evolution

Values of *λ <* 1 imply a component of variation in **y** that cannot be explained by phylogenetic dependence, conditional on a phylogenetic tree (topology and branch lengths) and a specified model of trait evolution. The default model of evolution in PMM is Brownian Motion (BM)— a continuous-time stochastic process characterised by a random walk of Gaussian distributed increments. The process branches at each node in the phylogeny with the property that trait variance increases linearly with time, meaning that shared branch lengths are proportional to the expected (co)variances of trait values among species. Thus, for BM, there is a direct and simple translation from phylogeny to covariance matrix (Ives 2018; Felsenstein 1985).

In the original presentation of phylogenetic generalised least squares (PGLS) (Grafen, 1989), a scalar multiple of the phylogenetic correlation matrix is taken to represent the total covariance among observations, Σ = *σ*^2^**C**, which assumes traits have evolved via BM on the tree with no additional variation. This assumption is equivalent to fixing 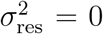 in (5). With subsequent developments, **C**, or equivalently the branch lengths of the corresponding phylogenetic tree, is transformed as a function of one or more additional parameters ***θ***, allowing different evolutionary models to be fit to the data (Symonds and Blomberg, 2014; Harmon et al., 2008). For a model with no additional random effects, this transformed phylogenetic correlation matrix **C**(***θ***), is then multiplied by the scalar variance parameter *σ*^2^ to define

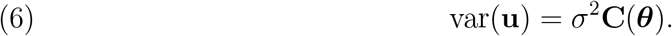

A common instance is when **C**(***θ***) is a weighted sum of **C** and an identity matrix **I** (equal in dimension to **C**) characterised by a single parameter, ***θ*** = *λ* (Pagel, 1999),

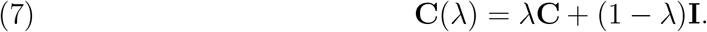

In this popular formulation of PGLS, the additive terms express the relative proportions, scaled by *λ*, of the total variance attributable to phylogenetic and residual effects. For the special case of a single observation per species (e.g., a species mean), (6) and (7) can be combined to obtain equivalence with the model given by (2) and (3) for 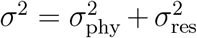 and 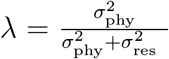 (also see Housworth et al. 2004; Cinar et al. 2022). This equivalence clarifies the relationships to other common methods: The basic PMM is equivalent to the original presentation of PGLS when 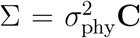 (i.e.,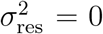) and to ordinary least squares when 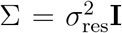 (i.e.,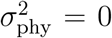) (Westoby et al., 2023; Blomberg et al., 2012). The phylogenetic threshold model of Felsenstein (2005) is equivalent to PMM with a binary response trait and probit link (Hadfield 2015; see section 3.0.5 for details).

In addition to (7), many alternative models for the evolution of continuous traits have been proposed, including those with directional trends, stabilising selection, or changes in evolutionary rates over time (Harmon et al., 2010; Butler and King, 2004; Blomberg et al., 2003; Pagel, 1999; Hansen, 1997). PMM supports several popular models (e.g., *δ, κ*, early burst (EB), Ornstein-Uhlenbeck (OU), accelerating-decelerating (ACDC)) by appropriate transformations to **C**, as in (6). This extends the estimation of phylogenetic random effects to a range of evolutionary models besides BM (see Tung Ho and Ané 2014; Blomberg et al. 2003), for which model selection techniques can be used to discriminate among alternative evolutionary hypotheses (Revell and Harmon, 2022; Goolsby et al., 2017). Unfortunately however, neither `MCMCglmm` nor `brms` currently provide options for estimating the parameters of evolutionary models other than BM.

#### 2.2.3 Limitations of single response models

PMM provides an elegant solution to the statistical problem of phylogenetic dependence in inter-species data. However, univariate implementations have important limitations that arise from treating one species trait, often arbitrarily, as the response variable and other traits as predictor variables. In particular, the validity of inferences can be compromised when predictor traits themselves contain phylogenetic signal (Westoby et al., 2023).

Suppose we observe a phenotypic correlation between two traits, **y**_1_ and **y**_2_, across a number of species and that both traits display phylogenetic signal; a common occurrence in empirical data sets (Adams and Collyer, 2019; Blomberg et al., 2003; Freckleton et al., 2002). Part of the covariance between **y**_1_ and **y**_2_ across species is due to a phylogenetically conserved relationship between these traits (e.g., phylogenetic niche conservatism, Losos 2008; Wiens et al. 2010), with the remainder due to causes that are independent of phylogeny (e.g., within-species trait covariance). If we model these data using a single-response model (1), treating **y**_2_ as a fixed predictor of **y**_1_ reduces this composite of correlations operating at the phylogenetic and non-phylogenetic levels to a single slope parameter. Not only are the separate (co)variance components not recoverable from the model, but phylogenetic signal in the predictor trait **y**_2_ is confounded with (phylogenetic components of) the residual covariance structure (Warton 2022; also see Wilson 2008; Rausher 1992 and Marques et al. 2022 for analogous issues in quantitative genetics and spatial statistics). In a worst-case scenario, these two correlation components may cancel each other out, leading to the erroneous conclusion that the traits in question are not meaningfully related (Westoby et al., 2023).

An alternative approach, which re-frames this analysis as a multivariate statistical problem, is to treat both traits **y**_1_ and **y**_2_ as response variables. That is, move all traits to the left-hand side of the model equation (9), into a stacked column-vector **y** = (**y**_**1**_, **y**_**2**_)^T^. This shift to a multivariate framework has several distinct benefits: 1) it avoids often arbitrary decisions about which trait should be considered the response variable and which the predictors, 2) it allows phylogenetic signal to be modelled in all traits simultaneously, 3) it permits a partitioning of trait correlations into phylogenetic and non-phylogenetic components, 4) it exploits correlations between response traits to improve precision in the prediction of missing and/or partially observed new data. We argue this approach is usually more appropriate and more meaningful for comparative analyses including multiple species traits (also see Section 4).

### 2.3 Multi-Response Phylogenetic Mixed Models (MR-PMM)

In MR-PMM, all species traits are modelled jointly as response variables, which means both the random effects **u** and residual errors **e** associated with each response must also be modelled jointly. For Gaussian response traits, this is achieved directly using multivariate normal distributions (but see Section 3.0.5 for non-Gaussian response types). Due to the compact form of the mathematical notation, the multi-response model is written identically to the single-response case,

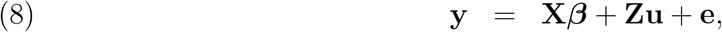

where 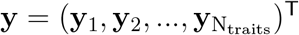 contains observations for all traits and all species, stacked in a single column vector. To illustrate, we represent the linear predictor for the bivariate case by placing each trait in a separate row,

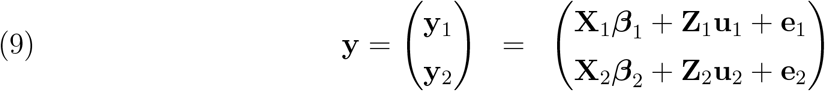

with

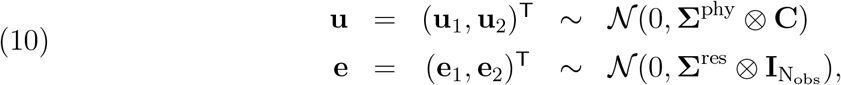

where **X**_1_***β***_1_ and **Z**_1_**u**_1_ are the linear predictors for the fixed and random effects, respectively, for response trait **y**_1_, and similarly for **y**_2_. The notation adopted in (9) emphasizes the joint distribution of effects for each response trait, where between-trait covariances across both phylogenetic **u** and residual **e** components are modeled with the trait covariance matrices

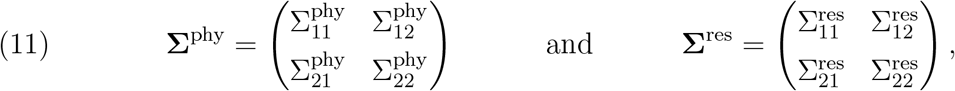

where the matrix elements 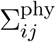 and 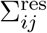 are scalar covariance components between traits *i* and *j*, representing the phylogenetic and residual components of covariation, respectively. The Kronecker product ⊗ is a type of matrix multiplication which, using the phylogenetic component as an example, can be written as

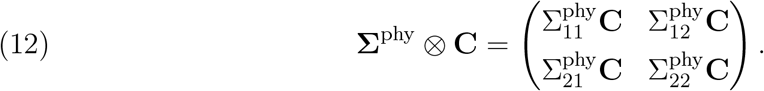

The Kronecker product has the effect of introducing phylogenetic dependence, via **C**, into the covariance structure of each trait and each pairwise trait relationship (see tutorial for fully worked examples). The total variance of **y** is the sum of the variances of the phylogenetic and residual components, transformed to the dimension of the multivariate data vector using the random effects design matrix **Z**,

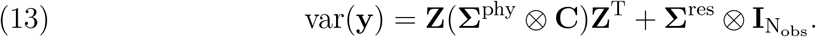

Despite the large dimension of var(**y**), the number of estimated (co)variance parameters in MR-PMM is N_trait_(N_trait_ + 1) (e.g., only 6 parameters for the bivariate case). For the general case, these parameters can be expressed in terms of the standard deviations *σ*_*i*_ and the correlations *ρ*_*ij*_ used to characterise the matrix elements of both 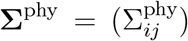 and 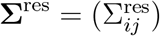 written as

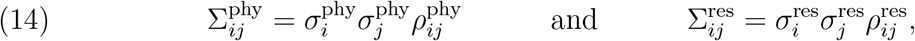

for *i, j* = 1, .., N_traits_.

#### 2.3.1 Implications

Moving from a SR to a MR model facilitates more refined inferences, improves prediction, and allows users to include all available data, not just complete cases. In terms of inference, the Kronecker products in (10) provides a means to include both phylogenetic and non-phylogenetic structures in the modelled (co)variance term for each trait and trait pair. Specifically, for the basic MR-PMM presented in (9) and (10), the estimated quantity 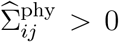 provides evidence for a component of correlation between traits *i* and *j* that is conserved over evolutionary time (i.e., a portion of trait covariance that is associated with phylogeny), while 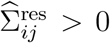 provides evidence for a component of correlation that is independent of phylogeny (see Figure 2 for an illustration). In contrast, the SR model fails to account for signal in ‘predictor’ traits, leading to a confound between phylogenetic and residual effects (Westoby et al. 2023, also see Section 2.2.3).

For predictive goals, MR-PMM is both more precise and more accurate than univariate linear models due to sharing of information across levels that accounts for phylogenetic relationships. Prediction to missing trait values leverages all data across traits and species, weighting their contributions by the estimated phylogenetic and non-phylogenetic covariances (see Section 5). Meaningful reductions in predictive variance compared to SR models may require that traits are at least moderately correlated (Riley et al., 2017; Jackson et al., 2011). This may apply even more so at the phylogenetic level, where estimates of covariance are generally more uncertain (see Tutorial). However, conserved trait correlations are not uncommon in nature (Westoby et al., 2023), with comparative studies often investigating hypotheses about coordination (e.g., trade-offs, co-selection) among species traits that display strong phylogenetic signal (e.g., Kelly et al. 2021; Sanchez-Martinez et al. 2020; Blomquist 2019). Indeed, recent work highlights the utility of MR-PMM for analysing function-valued traits such as performance curves, growth trajectories, or reaction norms (Pottier et al., 2024). Parameters that describe these functions (e.g., intercepts, slopes) are often correlated, and the strength and direction of these correlations may be biologically meaningful, providing opportunities to test hypotheses about the functional form of trait relationships (e.g., Pettersen et al. 2023; Kontopoulos et al. 2020).

Finally, the structure of the MR model allows us to flexibly include missing data. This means that users can retain observations where only a subset of the response traits have been observed, which improves inference (e.g., observations reporting values for just two response traits out of the complete set improve the estimation and decomposition of the (co)variances for these two traits). Including incomplete records usually means that more data (observations and species) can be included in analyses, especially as the number of included response variables increases and incomplete cases may comprise a substantial fraction of the data (see Tutorial).

## 3 MR-PMM - extensions to the basic model

In this section, we explore several useful extensions to the basic MR-PMM that expand the utility for comparative biology.

### 3.0.1 Multilevel models

Data for phylogenetic comparative analyses are typically compiled from multiple sources. This often yields multiple observations per species, with individual studies contributing observations on only a subset of focal traits for a subset of species. Mixed models like MR-PMM offer a general framework for dealing with these complex, cross-classified effect structures via multilevel models. Benefits include the ability to partition variation among multiple hierarchical levels (e.g., between-species, within-species, within-individual), account for variation in sampling effort, and fit models to data containing a mixture of record types (McNeish 2021; Nakagawa and Santos 2012, also see Figure 4 in Nakagawa et al. (2017) for a helpful visualization). Furthermore, accounting for multilevel structure may be necessary for valid inference of phylogenetic (co)variances, which are likely to be confounded with other sources of between-species variation (Cinar et al., 2022; Garamszegi and Møller, 2017).

Equation (9) can be extended to account for multilevel structure in the data by modelling additional random effect components in **u** that capture the sampling hierarchy. For example, to account for study effects, we include a random effect at the study level **u**^study^, which models variability between studies. To account for multiple observations per species, we include an additional, unstructured random effect at the species level **u**^ind^, which models variability between species that is independent of phylogeny. Each component term of **u** is associated with a (co)variance structure that specifies dependence within that component term (15). However, the effects of different component terms are assumed to be independent *a priori*,

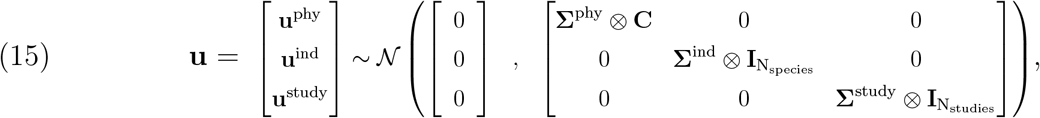

where 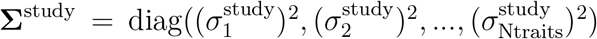. Modelling these additional components in **u** has important consequences for the way trait variation is partitioned, affecting the interpretation of model parameters. For example, the random effects specified in (15) represent a shift from modelling all non-phylogenetic sources of trait correlation in the residual error **e** as in (9), to modelling non-phylogenetic between-species trait correlation explicitly, via **u**^ind^, study effects via **u**^study^, and any remaining trait (co)variance as residual error in **e**. Residual correlation for such a model represents within-species trait correlation, which may require very large datasets to be robustly estimated (Zhou et al., 2022). In practice, this means residual errors are often treated as independent in multilevel MR-PMM (i.e., **Σ**^res^ is constrained to a diagonal matrix). For example, by setting `rescor = F and idh(trait):units in brms and MCMCglmm`, respectively (see tutorial).

Additional components could, of course, be added to **u** (15) to model other sources of covariance between observations, such as spatial or temporal effects (e.g. Markovski et al. 2023; Gomes et al. 2023; also see Freckleton and Jetz 2009). This flexibility of multi-level models makes MR-PMM well-suited for large-scale comparative studies including complex hierarchically structured data.

### 3.0.2 Meta-analyses

Data compiled for phylogenetic comparative analyses often contain a mixture of record types, including individual-level observations as well as species means, with or without associated sampling variances. In some instances, effect size measures such as test statistics or model parameters with associated errors are specified as response variables. Integrating data with heterogeneous errors casts MR-PMM as a meta-analysis in which observations are weighted according to their uncertainty. Multi-response meta-analyses have become increasingly common in clinical medical research over the past decade (Jackson et al., 2011; Riley et al., 2017), and are currently in focus for researchers in ecology and evolution (Pottier et al., 2024).

We emphasise that it is always preferable to fit models to observation-level data rather than statistical summaries of those data, such as means with associated sampling variances. In particular, models fit to species mean trait values neglect within-species variation, which can bias results (e.g., estimates of phylogenetic signal, Cinar et al. 2022) and ultimately obscure the relative contribution of biological and methodological drivers of the observed trait variation. However, meta-analytic implementations of MR-PMM may be desirable when summary statistics are the only available data for some species. In practice, observation-specific sampling variances 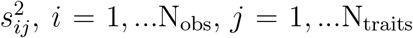 are often modelled as additive components of the residual variance,

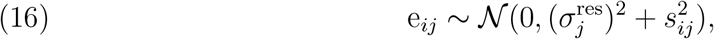

for example, using the `se()` and `mev()` arguments in `brms` and `MCMCglmm`, respectively. However, this approach is limited for data with a mixture of records, because individual-level observations are rarely reported with observational error, meaning only the sampling variances of species mean observations can be modelled. A straightforward approach is to weight each observation proportional to its reported sample size. This is implemented via the `weights` argument in `brms`, though currently unavailable in `MCMCglmm`. One workaround is to produce a vector of inverse model weights to be passed into the model as sampling variances (e.g., *w*^−1^ = 1*/n*). However, this leads to scale issues, where the sampling variance added to individual-level observations (*n* = 1) may be large relative to the residual error.

### 3.0.3 Distributional models

Distributional models allow separate predicition for each parameter of the response distribution, e.g., the mean and variance of a Gaussian distribution (also known as a location-scale model). Biologically, these models distinguish between the central tendency (location) and variability (scale) of a trait, both of which can be influenced by genetic, environmental, and evolutionary factors (O’Dea et al., 2022; Westneat et al., 2015). Multilevel distributional models that include random intercepts in both the mean and variance, for a common grouping variable, introduce covariance and heteroskedasticity (non-constant variance), respectively—so called double-hierarchical models.

To illustrate, consider the multilevel MR-PMM discussed in Section 3.0.1. The **u** terms in (15) are *location* random effects, because each level in the hierarchy (e.g., **u**^phy^) adds a normally distributed random intercept to the model for the *mean*, which captures covariance attributable to the grouping variable. However, the model remains homoskedastic across species—all observations, irrespective of their phylogenetic effects, share a common residual variance—unless the model for the residual variance also depends on the group structure.

To implement a group-structured variance model, we can specify

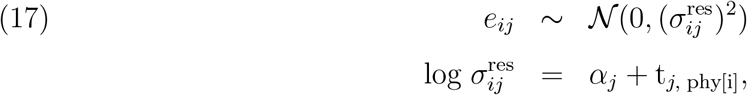

where, 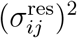 is the residual variance of the *i*^*th*^ observation for the *j*^*th*^ trait, *α*_*j*_ is the fixed intercept for the variance of the *j*^*th*^ trait, and t_*j*, phy[i]_ is a trait-specific random phylogenetic effect. To include the possibility of correlations between the random phylogenetic effects appearing in both the mean and variance, and thus share information across the distributional parameters, it is preferable to specify a joint prior distribution (Bürkner, 2019),

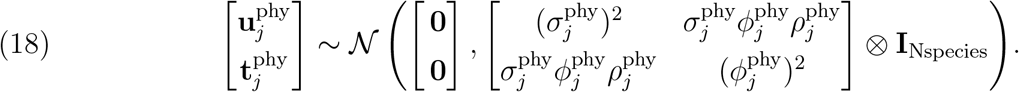

While distributional models are increasingly used to understand variation in intra-individual behavioural variability (Mitchell et al., 2021; O’Dea et al., 2022), models such as (18) appear to be severely underutilised in phylogenetic comparative analyses (Horvath et al., 2023; Cleasby and Nakagawa, 2011). Indeed, the mean and variance of biological traits are often positively correlated (a special case of heteroskedasticity), routinely threatening the assumption of homogeneous variances in common statistical tests. In some instances, this issue is addressed by the mean-variance relationship determined by the choice of statistical distribution and link function (e.g., Poisson with log link), or else using variance-stabilising transformations (e.g., log transforming response data). However, heteroskedasticity may remain in spite of these efforts, and this ‘variance of the variance’ may in fact correspond to interesting biological effects that we would like to model explicitly. As with our model for the mean, we may include an arbitrary number of predictors for the variance, including fixed effects (e.g., methodological or environmental covariates), and random effects that contain further hierarchical structure. For example, a reasonable assumption for many phylogenetic comparative analyses is that residual variances should differ between species and that variation in residual variance should itself contain phylogenetic structure (i.e., closely related species should show similar amounts of residual variability). Furthermore, estimating correlations between phylogenetic random effects (as in 18) provides a method to test for conserved mean-variance relationships within and between traits.

For studies attempting to understand drivers of phenotypic variability, explicitly partitioning trait variance across known hierarchical structures has the potential to be extremely informative (O’Dea et al., 2022; McNeish, 2021; Westneat et al., 2015). Distributional models also have a clear statistical motivation; heteroskedasticity violates, by definition, the assumption of homogeneous variances, leading to inefficient parameter estimation and biased errors, with potential consequences for inference and hypothesis tests.

### 3.0.4 Multivariate models of evolution

In recent years, there have been considerable advances in software implementation and efficient computation of multivariate evolutionary models (Bartoszek et al., 2023; Blomberg et al., 2020; Mitov et al., 2020; Clavel et al., 2015; Bartoszek et al., 2012). Models of evolution comprise both stochastic and deterministic components and are naturally expressed as stochastic differential equations where the rate of change of trait values over time depends on current values. Solutions (i.e., the distribution of trait values) are obtained by integrating these equations over the phylogeny. For many multivariate models (e.g., multivariate BM, OU, EB and ACDC), the trait distributions are Gaussian and therefore characterised by their means and covariances. In the multivariate BM case, the solutions are more easily obtained (though still present convergence issues even for the most sophisticated algorithms (Butler et al., 2023)) since the covariance is proportional to **C** (12) up to a re-scaling of the off-diagonal elements (13), equivalent to a simple branch-length transformation. For more complex models of evolution, the covariance matrix cannot generally be obtained via branch-length transformation, which appears to limit the scope of MR-PMM in accommodating alternative evolutionary models. However, in certain cases, analytic expressions for the total covariance have been derived. To illustrate we consider the most general multivariate OU model,

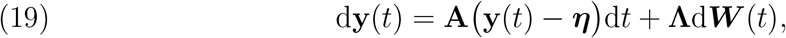

where, **y** is an N_traits_-vector of trait values, **A** is a N_traits_ *×* N_traits_ strength of selection (or rate of adaptation) matrix, ***η*** is an N_traits_-vector of trait optima values, and ***W*** is an N_traits_-dimensional Brownian process with diffusion matrix **Λ**. Viewed in discrete time as a vector autoregressive model, the differential equation (19) provides a rule to update trait values based on their current values together with a (Brownian) stochastic step. Each off-diagonal element of **A** models the deterministic effect of one trait value on the update of another trait, permitting a test of Granger causality—the notion that historical values of one trait improve the prediction of another (Shojaie and Fox, 2022). Although Granger causality provides a more direct inference of causation than a correlation coefficient, especially given **A** in (19) need not be symmetric, it is not necessarily more meaningful than the trait correlations that we can derive from **Λ**. Indeed, estimated correlations in the stochastic step may also capture causal relationships, just not those associated with adaptation toward trait optima.

For the MVOU model (19), the N_traits_ *×* N_traits_ trait covariance matrix between two species *i* and *j* at the tips is given by (Bartoszek et al. 2012, eq. B.3; also see Clavel et al. 2015),

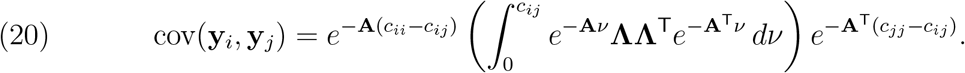

where the phylogeny enters the computation via the matrix elements of **C** = (*c*_*ij*_). As intimidating as it appears, (20) can be evaluated directly provided the matrix **A** admits an eigendecomposition (eq. A.1 in Bartoszek et al. 2012) which is likely to be satisfied under most circumstances (Mitov et al., 2020). In the limiting case that **A** goes to zero (i.e., no attraction to optima), (20) reduces to the standard multivariate BM where **Σ**^phy^ = **ΛΛ**^T^. Many common MVOU variants are characterised by strong constraints on the form of **A** which further simplify the evaluation of (20). For example, when attraction to optima is assumed to be equal and independent for all traits; i.e., when **A** is a multiple of the identity matrix (Goolsby et al., 2017; Tung Ho and Ané, 2014).

In principle, analytic computation of cov(**y**_*i*_, **y**_*j*_) extends MR-PMM to a broad class of multivariate Gaussian evolutionary models (e.g., for multivariate OU, it may be sufficient to use (20) in place of **Σ**^phy^ ⊗ **C** in (9)). However, options to implement models other than BM are yet to be integrated into popular software packages such as MCMCglmm and brms, despite being well developed in other modeling contexts (Bouckaert et al., 2019; Mitov et al., 2020; Clavel et al., 2015; Goolsby et al., 2017). A valuable direction for future work would therefore be to explore the repertoire of evolutionary models compatible with MR-PMM and integrate them into standard software; together with modern algorithms for computing tree-structured multivariate likelihoods (Mitov et al., 2020; Hassler et al., 2022; Bastide et al., 2021). Progress in this direction would yield a powerful modelling framework, capable of synthesising model selection among alternate multivariate evolutionary models with the generalisable multilevel framework of MR-PMM.

### 3.0.5 Non-Gaussian response traits

One advantage of MR-PMM compared to other methods (e.g., Goolsby et al. 2017; Clavel et al. 2015; Bartoszek et al. 2012) is the capacity to include both continuous and discrete response traits. To simultaneously include different trait types, a latent-variable formulation is used. This approach models between-trait covariance in the usual way, via a multivariate Gaussian distribution, while at the same time permitting trait-specific probability distributions and associated link functions (Hadfield, 2010). For traits *i* = 1, .., N_trait_, the latent-variable formulation is written

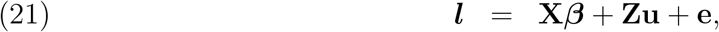

where 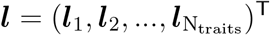 are latent variables associated with the observed traits and the corresponding (latent-trait) covariances are given by matrices **Σ**^phy^ and **Σ**^res^ which characterise the distributions of **u** and **e** as in (10); the latter are now ‘pseudo’ rather than true residuals introduced as observation-level random effects. A model for the observations **y**_*i*_ is given in terms of a trait-specific probability distribution *f*_*i*_ and link function *η*_*i*_

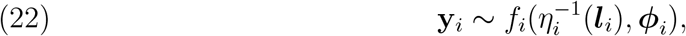

where ***ϕ***_*i*_ includes any distribution-specific parameters (if required). Depending on the choice of distribution *f*_*i*_, additional constraints may need to be imposed on associated variance terms to ensure parameter identifiability.

Variance partitioning to define an analogue of *λ* (5) is possible for the latent-variable models given by (21) and (22), but is more nuanced because there is a distinction between latent- and observation-scale partitioning where distribution-specific variances must be taken into account (Nakagawa et al., 2017). Variance terms are usually more easily interpreted when they are expressed on the scale of trait observations rather than that of the latent variables (de Villemereuil, 2018). For multi-response models involving two or more distribution types, there are generally no closed-form solutions for the observation-level variances and associated statistics, however numerical methods are readily applied given point estimates or a set of posterior samples of the model parameters (de Villemereuil et al., 2016). A further complication for interpretation, however, is that the non-linear transformations required to transform covariances from the latent to the observation scale can change the bounds of the correlation coefficient. For example, the minimum correlation of two exponentiated random normal variables is approximately −0.37 and the correlation bounds of a random normal and an exponentiated random normal variable are approximately *±*0.76.

An alternative to specifying trait-specific probability distributions, is to model discrete or constrained trait observations as censored measurements of continuous latent variables. This idea generalises the threshold model (Albert and Chib, 1993) where the absence of non-linear link functions means that correlations between the continuous variables, both observed and imputed, can be directly interpreted on the scale of observed data (Clark et al., 2017). Implementation of this approach requires sampling from truncated multivariate normal distributions for which recently developed computational methods have substantially increased efficiency, especially for high-dimensional problems which were previously intractable (Zhang et al., 2023).

The software packages `MCMCglmm` and `brms` incorporate non-Gaussian traits using the approach given by (21) and (22), whereas `BEAST` implements the generalised probit model using the new efficient sampling approaches.

### 3.0.6 Considerations for fixed effects

There are two common reasons for including fixed effects in a MR-PMM. The first is to account for sampling biases, such as differences in data-collection methods or laboratory protocols. For example, in the field of plant hydraulics, physiological traits related to xylem vulnerability are often measured using different techniques that each have distinct assumptions and sources of error (Jansen et al., 2015). In such situations, it is advisable to account for measurement methodology with a fixed effect covariate. The second reason is to explore relationships between responses conditional on covariates, typically to evaluate hypotheses about the mechanistic basis of trait correlations. For example, Sanchez-Martinez et al. (2020) used MR-PMM to partition correlations between a range of plant hydraulic traits into phylogenetic and non-phylogenetic contributions, then re-fit models with relevant climate covariates included as fixed effects. Whether or not trait correlations remained after accounting for climate effects was used to evaluate hypotheses about trade-offs and integration between these traits in the evolution of plant hydraulic systems. Equivalent procedures are used to control for environmental variation in multivariate species co-occurrence models (Ovaskainen et al., 2017; Warton et al., 2015). However, in MR-PMM, a preferable approach will often be to treat biological covariates as response traits and assess conditional dependencies via partial correlations.

### 3.0.7 Partial correlations

The practice of assessing conditional relationships between traits (i.e., relationships after controlling for relevant covariates) is well-established for univariate models. For example, it is common for researchers to use multiple regression to estimate partial regression coefficients for each fixed effect predictor. In MR-PMM, we estimate conditional relationships from a joint model, where both focal traits and covariates are modelled as response variables. This is preferable because traits commonly treated as covariates, such as those characterising the climatic or environmental niche of species, often display strong phylogenetic signal (Liu et al., 2020; Kubota et al., 2017; Peixoto et al., 2017; Hof et al., 2010; Kozak and Wiens, 2010; Smith and Beaulieu, 2009; Hawkins et al., 2007). Thus, treating covariates as response variables can avoid potential confounds between fixed and phylogenetic effects (see section 2.2.3). Partial correlations between response traits in MR-PMM are derived from elements of the inverse trait-covariance matrices, called precision matrices **Ω** = (Ω_*ij*_); that is,

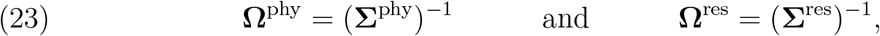

for which the corresponding partial correlation coefficients are 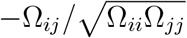.

The estimation of partial correlations via precision matrices casts MR-PMM in the framework of Gaussian graphical models (Popovic et al., 2019; Epskamp et al., 2018; Yuan and Lin, 2007; Magwene, 2001), which have great potential for clarifying assumptions of phylogenetic comparative methods (Uyeda et al., 2018). Elements of these precision matrices relate directly to the existence of edges in a graphical causal network (Figure 1). For example, suppose we obtain data on three co-varying species traits *x, y* and *z*. In the precision view (Figure 1 right panel), the absence of an edge between *x* and *y* signifies conditional independence: that *x* and *y* are independent, given *z*. This does not imply that the correlation between *x* and *y* is zero (Figure 1 left panel), only that the partial correlation between these traits is zero. Precision matrices are often sparser than their corresponding covariance matrices (i.e., contain more off-diagonal elements that are effectively zero), which focuses our inference on a reduced set of candidate causal relationships. For example, Halliwell et al. (2024) used MR-PMM to estimate partial correlations between social behaviour (binary response) and a range of species traits and environmental niche variables in a global sample of 1696 lizard species. The authors used partial correlations to disentangle direct climate effects on the evolution of social behaviour from indirect effects driven by adaptations to climate (i.e., traits correlated with climate) that go on to promote the evolution of social behaviour.

**Figure 1:**
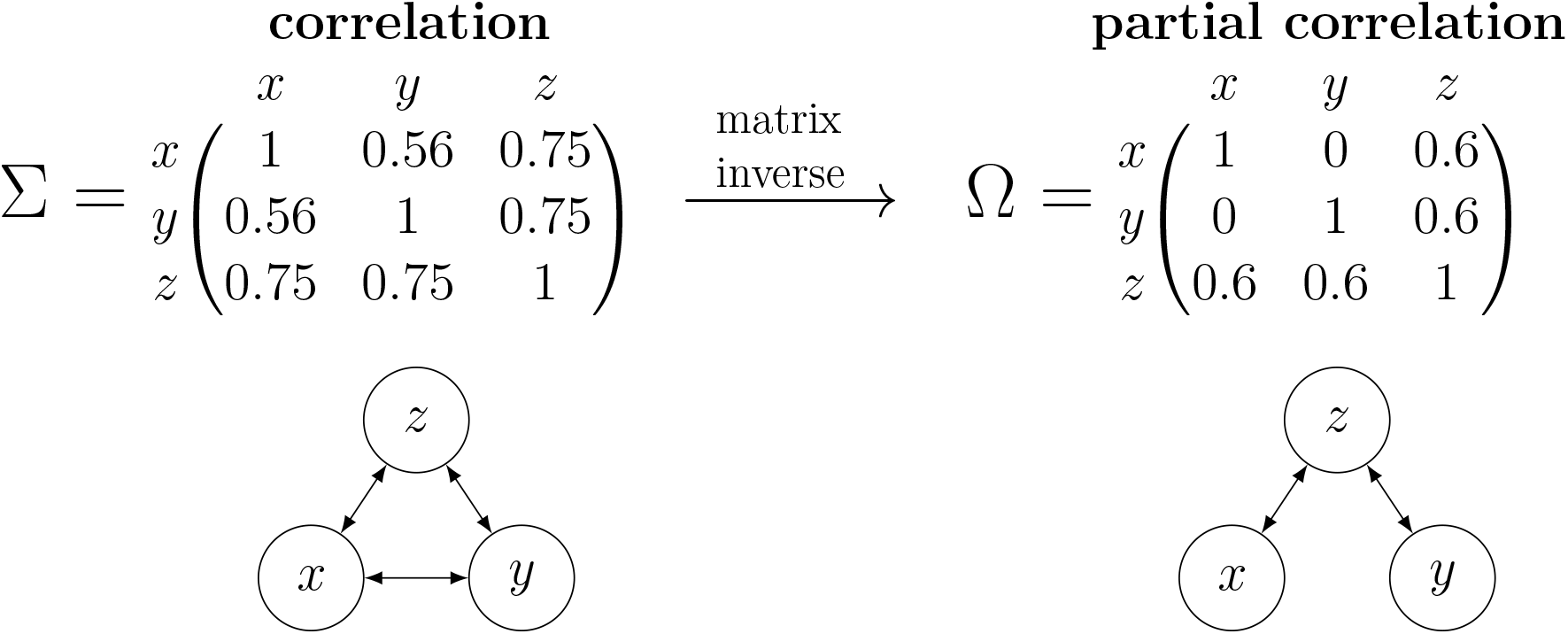
Correlation (Σ) and partial correlation (Ω = Σ^−1^) matrices between species traits *x, y* and *z*. The undirected network graphs below provide a qualitative representation of trait relationships, where the absence of an edge between two traits corresponds to a zero off-diagonal element in the matrix above. To evaluate whether the relationship between *x* and *y* can be explained by a covariate *z*, we compute partial correlations from the precision matrix of trait covariances, which quantifies the relationship between *x* and *y* when controlling for *z*. In this example, *x* and *y* are strongly correlated, however this relationship is fully explained by a shared response to the covariate *z*, i.e., *x* and *y* are independent, conditional on *z*.

Outside of controlled experiments, there is always the strong possibility that partial correlations could be explained by missing variables. Nonetheless, precision matrices are a powerful tool for generating candidate causal hypotheses from observational data (Popovic et al. 2019; Pearl 1995; also see tutorial), and are readily obtained from a fitted MR-PMM.

## 4 Interpretation

The biological interpretation of phylogenetic models and their utility for addressing specific ecological and evolutionary hypotheses has been a subject of lively debate (Freckleton et al., 2002; Westoby et al., 1995; Björklund, 1997; Harvey et al., 1995). Reflecting on this constructive body of work, we believe MR-PMM emerges as a flexible, pluralistic framework for analyses of multiple species traits. However, consensus on interpretation is critical to realise the full, operational potential of the method.

### 4.1 Phylogenetic (co)variances

Phylogenetic (co)variances estimate the conserved component of species phenotypes, given a model of evolution and a hypothesis of the phylogenetic relationship between taxa. The extent to which these conserved effects result from stochastic processes, or reflect constraints and adaptive responses to selection, will vary for different traits, between clades, and across phylogenetic scales. These forces are also likely to fluctuate throughout the course of evolutionary history, complicating the interpretation of trait covariance (Revell et al., 2008; Losos, 2008, 2011). As Housworth et al. (2004), point out “[PMM] envisions phenotypic evolution as being the result of a complex of forces including some that are retained over long periods of time, forming patterns in trait variation that reflect the underlying phylogenetic structure”. Expressed another way, the phylogenetic component of trait (co)variance in PMM models variation attributable to the phylogenetic relationships among taxa. Examples include variation arising from genetic differences between clades that have accumulated over evolutionary time, as well as non-genetic (e.g., spatial, environmental or developmental) effects that are phylogenetically structured for one reason or another.

### 4.2 Pattern versus process

There are inherent limitations to the inferences we can make about evolutionary processes from data observed only in the present day. Indeed, many have argued that the interpretation of phylogenetic comparative analyses should be limited to patterns of variation, rather than the explicit processes that generated them (Revell et al., 2008; Losos, 2008, 2011; Ives, 2018). Others have suggested that integration of fossil evidence, path analyses, ancestral state reconstruction, and simulation studies may extend our epistemic reach to hypothesis tests of the processes generating observable variation (Thorson and van der Bijl, 2023; Uyeda et al., 2018; Quental and Marshall, 2010; Slater et al., 2012; Uyeda and Harmon, 2014).

Because phylogenetic and non-phylogenetic correlations can arise from multiple distinct causal processes, we argue for caution around mechanistic interpretations. Two traits may be phylogenetically correlated if, for example, conserved genes underlying a set of traits show linkage or pleiotropy that constrains the evolutionary potential for certain trait combinations, or because traits form part of a coordinated life history strategy that involves phylogenetic niche conservatism (Westoby et al., 2023; Wiens et al., 2010). Phylogenetic correlations may also reflect indirect relationships between traits, due to similar selective pressures acting on multiple traits independently throughout evolutionary history. As above, such hypotheses should not be considered mutually exclusive (Losos, 2011; Revell et al., 2008). Deriving conditional dependencies can help evaluate evidence for alternative mechanistic hypotheses in MR-PMM (see Section 3.0.7). Ultimately, however, it is rarely the pleasure of a comparative biologist to declare causation. Rather, to uncover meaningful patterns of variation, test predictions from theory, and generate hypotheses for future comparative and experimental research.

### 4.3 Visualisations

To compare and contrast phylogenetic and non-phylogenetic (co)variance components, and clarify their biological interpretation, we simulated data from a simple MR-PMM for two Gaussian response traits y_1_ and y_2_, as in (9). Figure 2 shows scatter plots of simulated data together with the tree used to derive the phylogenetic correlation matrix *C*. The ellipses within the scatter plot provide a visual representation of the simulation conditions for each panel. A heat map of trait data plotted against the phylogeny highlights the distinct signature each source of covariance leaves in the data (see Figure 2 caption for details).

Visualizations of simulated data are powerful heuristics for biologists, as they provide tangible representations of the abstract covariance structures we aim to partition with MR-PMM. For more technical coverage of simulating multivariate data containing phylogenetic and non-phylogenetic covariance structure, we include detailed examples and R code in the tutorial.

### 4.4 Relevance for biological hypotheses

We see the flexible framework and nuanced interpretation of MR-PMM as having at least four strong usage cases for biologists: 1) to provide a more meaningful decomposition of trait (co)variances, including phylogenetic and non-phylogenetic components, among both Gaussian and non-Gaussian response traits; 2) to test for partial correlations, consistent with functional relationships between species traits, in a rigorous phylogenetic framework; 3) to test theory and generate new hypotheses about the ecological and evolutionary drivers of trait variation; 4) to predict species vulnerability to environmental change based on phylogenetically structured trait-trait and trait-environment relationships. With an emerging synthesis of evolutionary, ecological and genomic approaches to comparative analyses (James et al., 2021; Smith et al., 2020), we concur with recent work recognizing the considerable potential of MR-PMM to advance our understanding of trait evolution, niche conservatism, and community assembly at numerous scales (Pottier et al. 2024; Westoby et al. 2023; Gallinat and Pearse 2021, also see Abrego et al. 2020).

## 5 Prediction

While the estimation of trait covariance, rather than trait prediction *per se*, is the focus of MR-PMM, the estimated covariance structure can be fully exploited for predictive goals. This approach improves on ordinary multiple regression for prediction as it makes use of the phylogenetic structure in the predictors. The predictive distribution of a fitted model is also useful for model validation, such as posterior-predictive checks, or model comparison using predictive assessment such as cross-validation. We now review conceptual aspects of predictive distributions in MR-PMM, and refer the reader to the accompanying tutorial for worked examples of how these predictive methods can be implemented in R.

### 5.1 Predicting to new or missing observations

Missing data is a perennial problem in comparative biology (Freckleton, 2009; Nakagawa and Freckleton, 2008). For many traits of interest, data collection remains too expensive, technically demanding, or logistically challenging to keep pace with the expansion of species phylogenies and functional trait databases. To illustrate, we consider another example from the field of plant hydraulic ecophysiology. Physiological resistance to xylem cavitation is central to water stress tolerance in plants and thus a key metric for evaluating species and ecosystem vulnerability to climate change (Brodribb et al., 2020; Choat et al., 2012). While new methods allow direct quantification of cavitation resistance in diverse species (e.g., *P*_50_, the water potential at which 50% of xylem conductivity is lost to cavitation (Brodribb et al., 2016, 2017)), observations require specialised equipment, technical proficiency, and are time consuming to obtain. However, because cavitation resistance forms part of a coordinated growth strategy involving conserved trait networks (Sanchez-Martinez et al., 2020; Skelton et al., 2021; Liu et al., 2024), phylogenetically structured trait correlations are likely to enhance our ability to predict this crucial, yet labor-intensive, trait where sufficient data on correlated traits are available. Recent work highlights strong correlations between xylem anatomical traits and *P*_50_ (Lens et al., 2022), suggesting functional indicator traits, some of which are cheaply and easily measured at scale (e.g., petiole XLA, Blackman et al. 2024). An MR-PMM specifying *P*_50_ (poor species coverage), functional traits (moderate species coverage), and environmental indices (good species coverage) as joint response variables could leverage all available data, phylogenetic relationships, as well as trait-trait and trait-environment correlations to impute missing *P*_50_ values for unobserved species.

In this multivariate mixed modelling context, we emphasise that prediction is both more precise and more accurate, relative to univariate linear models, because information is shared across species in a way that respects evolutionary history. The MR-PMM predictive model for missing trait values learns from all observed data across traits and species, weighting their predictive contributions according to the estimated phylogenetic and non-phylogenetic covariances, given the chosen model of evolution.

### 5.2 Model validation using posterior predictive checks

In Bayesian settings, posterior predictive checks use simulated data from the fitted model to test both the adequacy of the fit to the data and the plausibility of the model predictions (Gelman et al., 2013). These checks are typically visual plots that rely on qualitative assessments (Gabry et al., 2019). To test for adequacy of fit, one option is to superpose the observed data onto a plot of the distribution of the simulated data. For a PMM, this type of check could be performed using a separate plot for each trait type with the observed data point for each species plotted on top of a 5 point summary of the predicted distribution (see Figure 3 for an example). For assessment of model plausibility, the current state of knowledge should be used to evaluate model predictions within ecologically plausible but perhaps unobserved ranges of the included covariates. For example, do predictions for ***y***_1_ make sense across the entire range of plausible ***y***_2_ values for a given set of taxa and/or region of interest?

**Figure 2:**
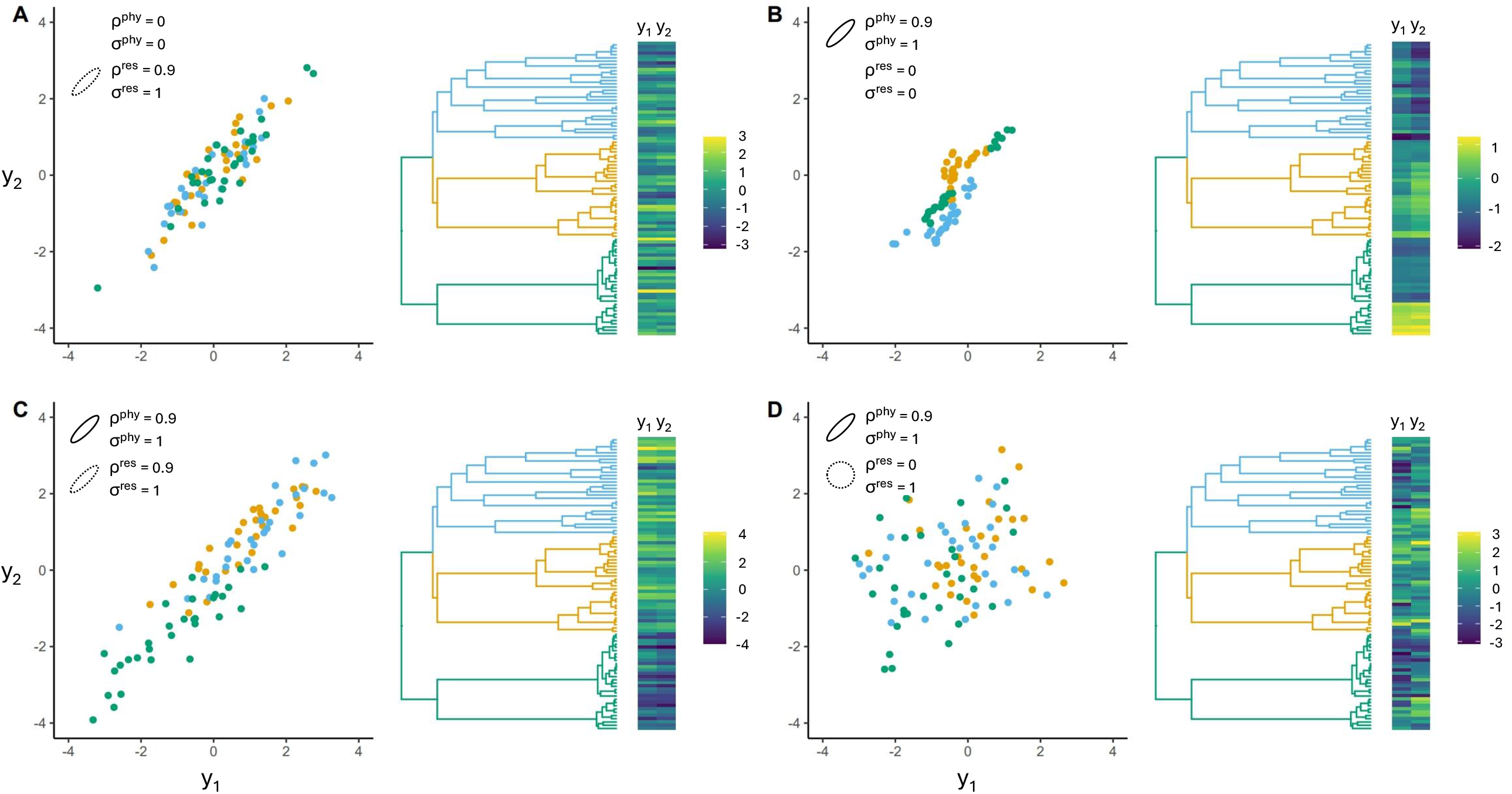
Bivariate trait data (y_1_, y_2_) simulated from a basic MR-PMM (9) containing different levels of phylogenetic (*ρ*^phy^) and residual (*ρ*^res^) correlation (**A**-**D**). Simulation conditions for each panel are inset at the top left of each scatterplot. Data are plotted in scatterplots, as well as heatmaps arranged against the generating phylogeny. In **A**, y_1_ and y_2_ have no phylogenetic signal (*σ*^phy^ = 0), but a strong positive residual correlation (*ρ*^phy^ = 0, *ρ*^res^ = 0.9). Clades overlap completely in the scatter plot and the heatmap shows bands of colour across y1 and y2 that appear random with respect to phylogeny. **B** shows the opposing situation, y_1_ and y_2_ are positively correlated but entirely with respect to phylogeny (*ρ*^phy^ = 0.9, *ρ*^res^ = 0) with no residual variation in either trait (*σ*^res^ = 0). The scatter plot shows clearly distinguishable clades and a tendency for both within- and between-clade correlation. The extent of between-clade correlation (the tendency for clades to arrange along a positive slope) depends on the topology of the tree, with deep splits promoting separation of clades along the major axis of co-variation. The heatmap shows phylogenetic structure weakening across clades from green, to orange to blue as the topology becomes more deeply nested, i.e., as subclades become less clearly separated in evolutionary time. In **C**, y_1_ and y_2_ have equal phylogenetic and residual variances, and a strong positive correlation operating on both levels. This scenario, where phylogenetic and residual correlations are similar in sign and magnitude, is likely to be common for many biological traits. In **D**, both traits contain phylogenetic and residual variance, but correlation is only present on the phylogenetic level. This shows how easily conserved correlations are obscured when residual sources of variation contribute substantially to trait variance. Notably, **D** represents a set of conditions for which SR-PMM, such as PGLS, will typically fail to detect a significant association between y_1_ and y_2_ (Westoby et al., 2023).

**Figure 3:**
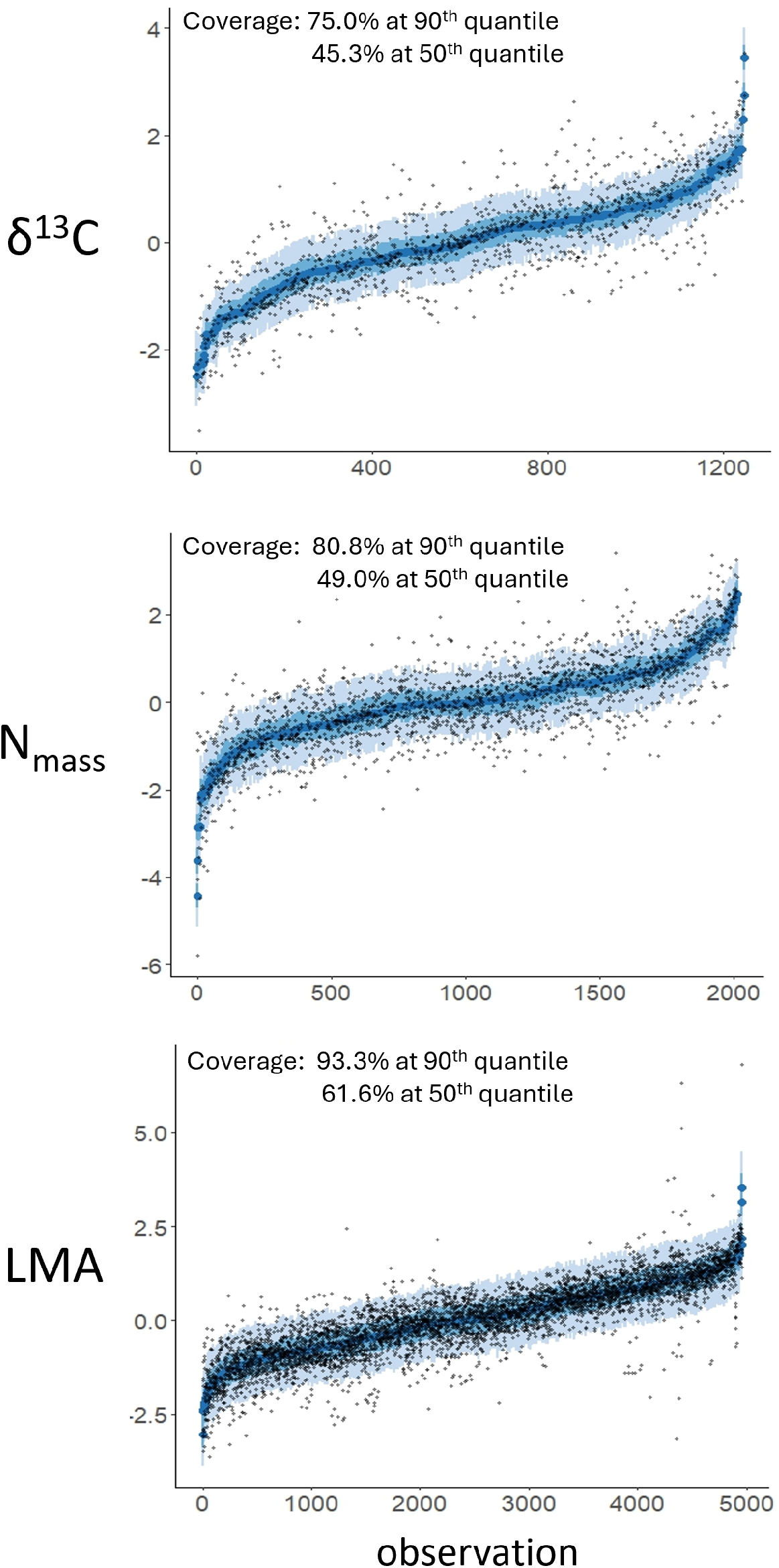
Posterior predictive checks from a fitted MR-PMM of leaf traits across 457 species of Eucalyptus. For each trait, black points represent observed values, while five-point summaries show the median (dark blue points), 0.5 CI (blue bars), and 0.95 CI (light blue bars) of the posterior predictive distribution for each observation. Predictions are made by conditioning on all random effects, with observations ordered by the predictive mean.

**Figure 4:**
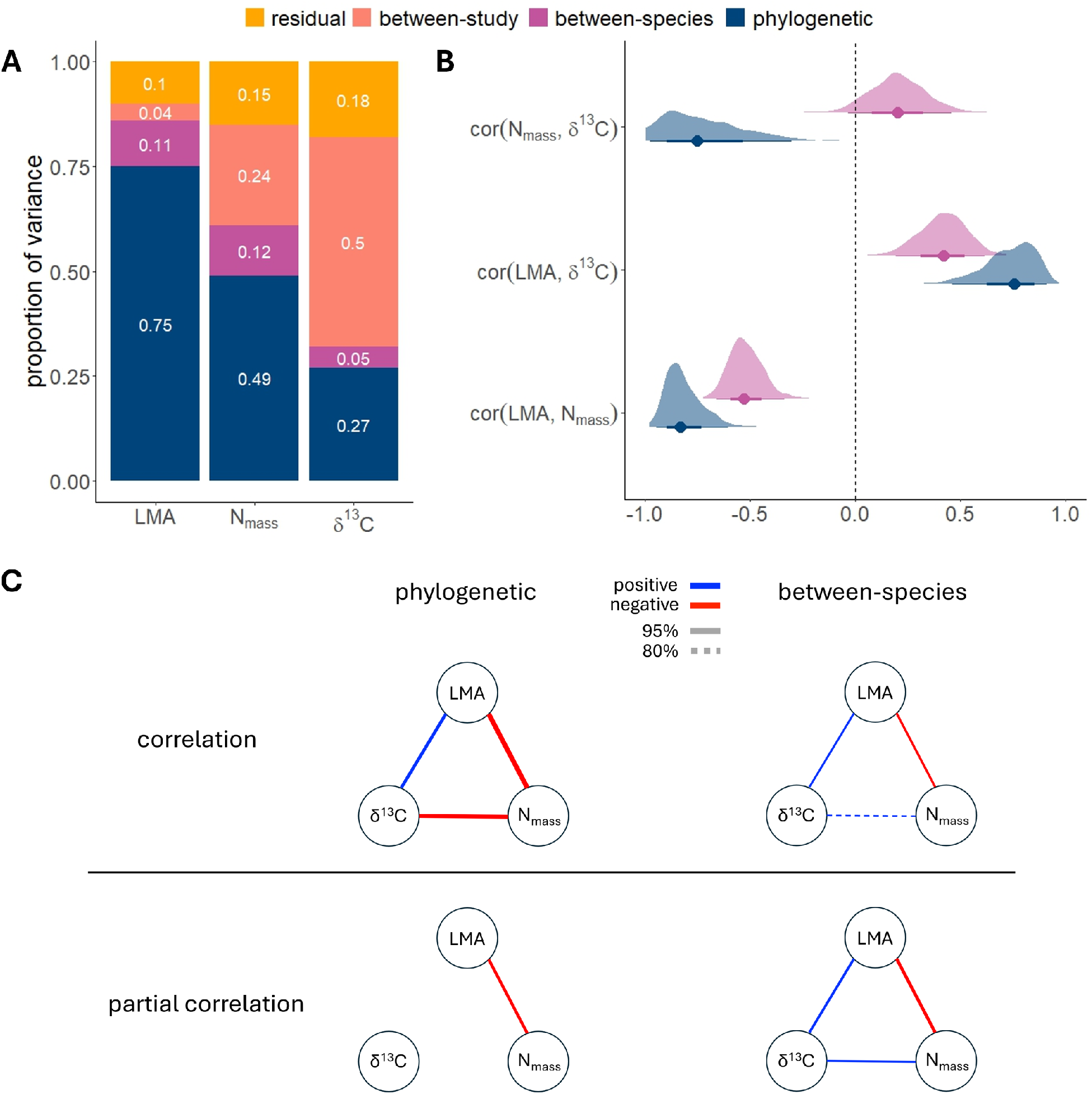
Results from an MR-PMM of leaf traits across 457 species of Eucalyptus. **A** variance decomposition reveals large differences between response traits in phylogenetic signal, as well as the relative contribution of different sources of variance; **B** phylogenetic and non-phylogenetic between-species correlation coefficient estimates. Points represent the posterior median for each estimate, with 50% and 95% CIs represented by heavy and light wicks respectively; **C** correlations and partial correlations between traits represented as network graphs. Edge widths are proportional to the posterior median of each coefficient estimate, with linetype indicating significance at the 95% (solid) and 80% (dashed) CI.

### 5.3 Predictive assessment using cross validation

Cross validation is the use of data splitting to estimate the predictive performance of one or more statistical models, usually for the purpose of model comparison, validation, or selection (Yates et al., 2022). Model selection is used when discrete decisions must be made about model structure. For example, whether or not to include various fixed effects (i.e., variable selection) or the choice of probability distribution (e.g., Poisson or negative binomial for count data). Predictive assessment tools such as cross validation are also useful to quantify or simply visualise how well a model can predict to new data (e.g., to new taxon-trait pairs in a PMM) which is distinct from typical assessments of model adequacy which concern prediction of data to which a model was fit.

Cross validation works by fitting each model to a subset of the available data and then assessing the models’ predictive capacities on the remaining data. The splitting procedure is systematically iterated to select different test data and the overall predictive performance is summarised as a cross validation score (Arlot and Celisse, 2010). When the measure of predictive performance is the log likelihood of the test data then the predictive assessment is said to be information theoretic. Information criteria such as Akaike’s Information Criteria (AIC, Akaike, 1973), or for Bayesian analyses the widely applicable information criteria (WAIC, Watanabe, 2010), approximate predictive log likelihood without data splitting by adding a bias correction to the log likelihood of the full data which requires only a single fit for each model. Information criteria are therefore faster to compute than cross-validation scores, however the latter are often preferred as they are less sensitive to violations of model assumptions and are readily combined with techniques to mitigate overfitting (Yates et al., 2021). For Bayesian MR-PMM estimated using Monte Carlo sampling, model fitting is often too slow to permit the use of ordinary cross validation, however recently developed approximate methods provide a rapidly computed and accurate alternative (Vehtari et al., 2017; Bürkner et al., 2021).

## 6 Example Analysis - leaf traits in Eucalyptus

To demonstrate potential applications of MR-PMM, we present an example analysis using data on leaf traits for 457 species of Eucalyptus from the AusTraits database (Falster et al., 2021). Our intentions for this analysis were to 1) decompose the variance in each trait and evaluate phylogenetic signal; 2) estimate phylogenetic and non-phylogenetic between-species trait correlations while accounting for multilevel structure in the data; and 3) evaluate partial correlations. Full details of this analysis, as well as additional examples exploring other model extensions with this dataset, are provided in the tutorial.

### 6.1 Methods

We derived the phylogenetic correlation matrix **C** from the maximum likelihood time-calibrated Eucalypt phylogeny “ML1” presented in Thornhill (2019). We focused on three target leaf traits: leaf mass per unit area (LMA); nitrogen content per dry mass of leaf tissue (**N**_mass_); and the ratio of carbon isotopes 12 and 13 in leaf tissue (*δ* ^13^**C**). All responses were log-transformed, zero-centred and scaled to unit variance prior to analysis. The random effects were specified as in (15), to estimate phylogenetic and non-phylogenetic between-species trait correlations while accounting for study effects.

For model validation, we performed posterior predictive checks. The proportion of the data falling in the predictive intervals was close to the nominal quantiles, indicating that the model is adequately calibrated (although the proportions for *δ* ^13^**C** were a little low), and visual inspection of the plotted distributions verified the capacity of the model to generate plausible data (Figure 3). The lower coverage of the posterior predictive intervals for *δ* ^13^**C** may indicate that the model is missing relevant covariates. Indeed, between-study effects accounted for the largest proportion of variance in *δ* ^13^**C**, suggesting methodological differences between studies or local environmental factors may be a contributing factor to poor predictive performance. Further, the chosen probability distribution (Gaussian), or the log-transformation, may be less well suited to this trait. For predictive assessment using approximate LOO-CV (Bürkner et al., 2021), see the tutorial.

### 6.2 Results

Phylogenetic signal, and hence the tendency for similar values among closely related species (Figure 5), varied considerably between traits (Figure 4**A**, posterior median for 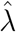: LMA = 0.75, **N**_mass_ = 0.49, *δ* ^13^**C** = 0.27).

**Figure 5:**
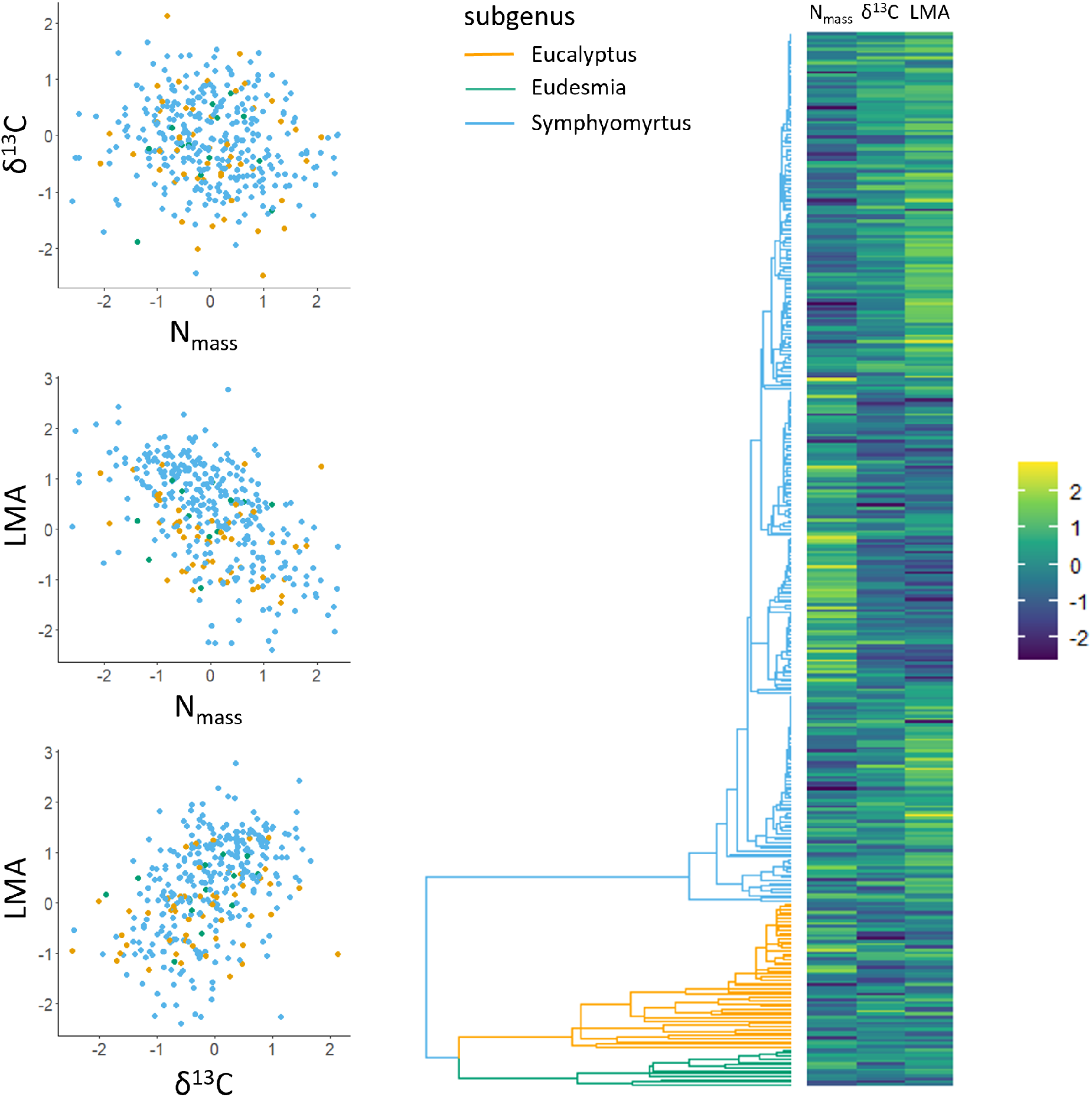
Scatter-plots (left) and heat-maps (right) of three leaf traits across 361 species of Eucalyptus (data filtered to complete cases). Trait values have been log-transformed and scaled. For heat maps (right), trait values are aligned with the corresponding species in the phylogeny (centre). For *δ* ^13^**C** and **N**_mass_ (top left), opposing phylogenetic and non-phylogenetic correlations reported by the model (Figure 4) are obscured at the level of species phenotypes.

The traits selected for this analysis are tightly linked to resource-use strategies and leaf economics (Reich, 2014; Wright et al., 2004). For example, Prieto et al. (2018) found that plant species in a Mediterranean woodland showing more resource acquisitive strategies (low LMA and high **N**_mass_), were associated with lower water use efficiency (low *δ* ^13^**C**). This is consistent with predictions from leaf economic theory (Wright et al., 2004), that LMA, *δ* ^13^**C** and **N**_mass_ represent components of a coordinated life history strategy which trade-off predictably as different regions of niche space are explored during species diversification. These co-evolutionary relationships are clearly supported by the fitted model (Figure 4**B**), which reports a positive phylogenetic correlation between LMA and *δ* ^13^**C**, and negative phylogenetic correlations between these traits and **N**_mass_. Thus, in line with theory and empirical observations from other plant groups, these traits appear to co-vary predictably over evolutionary time in Eucalyptus.

A notable result is that for **N**_mass_ and *δ* ^13^**C**, the non-phylogenetic correlation is positive while the phylogenetic correlation is negative (Figure 4**B**). This situation highlights an important strength of MR-PMM; the capacity to disentangle trait relationships operating at different levels in the model hierarchy. In line with simulations, consistent correlations across levels produce clear trends in inter-species data, while opposing correlations tend to obscure these relationships (Figure 5; Figure 2D; see tutorial).

Finally, while all non-phylogenetic trait relationships were retained as partial correlations, only the negative relationship between LMA and **N**_mass_ was retained at the phylogenetic level (Figure 4**C**). LMA and **N**_mass_ also showed the strongest relationships overall (Figure 4**B**), suggesting a deeply conserved functional integration between these traits.

## 7 Extended Topics

The following section explores technical, yet important aspects of fitting MR-PMMs. These topics are particularly relevant in the case of high-dimensional and/or sparse trait data, where issues such as prolonged fitting time, model convergence difficulties, or parameter identifiability require the application of specialised computational methodologies.

### 7.1 Priors for multivariate normal distributions

As discussed in Section 3.0.7, trait covariance matrices and their inverses are often targets for inference; for example, to estimate conditional dependencies between traits or compute phylogenetic signal. However, it can be challenging to estimate these matrices as they generally have a large number of parameters; there are 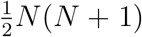 parameters in each of **Σ**^phy^ and **Σ**^ind^, where *N* = *N*_trait_ denotes the number of traits throughout this section. In the MR-PMM framework, the covariance matrices **Σ**^phy^ and **Σ**^ind^ each parameterise a multivariate normal distribution which are priors for the vectors **u**_phy_ and **u**_ind_, respectively (see (15)). In a fully Bayesian setting, it is also necessary to specify priors for the covariance matrices themselves, for which there are many different options to choose from depending on the way the model is parameterized. The choice of prior impacts on the model-fitting strategy and efficiency, as well as the degree and type of shrinkage.

The classical choices of parameterisation for the multivariate normal are the covariance and precision forms. To each of these choices, there is a conjugate prior, the inverse Wishart and the Wishart distributions, respectively, characterized by a target matrix and degree of belief parameter. Conjugate priors are an algebraically convenient choice because they provide analytic solutions for certain steps of the model fitting which can greatly reduce estimation time in a Gibbs sampling approach (the covariance form with inverse Wishart prior is implemented in MCMCglmm). Departing from conjugacy, regularising priors such as the zero-centred Laplacian (two-sided exponential) and Gaussian distributions can be used to ‘shrink’ the off-diagonal elements of the covariance or precision matrix. In particular, the use of the Laplacian prior for the precision matrix, called the *graphical lasso*, is able to shrink matrix elements all the way to zero, generating a sparse network of conditional dependencies. Further examples of sparsity-inducing priors include the adaptive lasso (Zou, 2006) and the horseshoe (Carvalho et al., 2009) which contain global terms to control the total amount of shrinkage and local terms to flexibly allocate shrinkage on a per-parameter basis. These types of sparsity-inducing priors have been used extensively for the general problem of variable selection, and hold enormous potential for inferring trait relationships in the MR-PMM setting, especially when data are sparse relative to the total number of candidate correlations.

A shortcoming of specifying priors for the covariance (or the precision) matrix is that the variances and correlations (or the precisions and partial correlations) are not treated independently. For example, we may wish to apply shrinkage to the correlation coefficients while maintaining an uninformative prior on the variances, or vice versa. To address this issue, the covariance matrix can be decomposed as **Σ** = **SRS**^T^ where **S** = diag(*σ*_1_, *σ*_2_, …, *σ*_*N*_) is the diagonal matrix of standard deviations and **R** is a correlation matrix (Barnard et al., 2000). This decomposition allows priors for the standard deviations to be specified separately from those of the correlation matrix. The matrix **R** can be further factored into the form **R** = **LL**^T^ where **L** is a lower-triangular matrix called the Cholesky factor. The latter parameterisation is implemented in brms where the default priors are half Student-t for the standard deviations and the Lewandowski-Kurowicka-Joe (LKJ, Lewandowski et al. (2009)) prior for **L**. The LKJ prior has a single parameter *η* which for *η* = 1 specifies a uniform distribution in the space of correlation matrices and for *η >* 1 regularises estimates toward the identity where each correlation coefficient *ρ*_*ij*_ has marginal density Beta 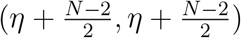.

Use of the Cholesky factor is computationally advantageous for multivariate normal models, as it simplifies evaluation of the likelihood and allows an unconstrained parameterisation of **R** in terms of correlation angles via a certain geometric representation (Forrester and Zhang, 2020). The latter are efficient to sample and allow rapid evaluation of the LKJ prior while guaranteeing the validity of the corresponding correlation matrix.

### 7.2 Latent factor methods for dimension reduction

Although regularising priors can reduce the effective number of parameters for an estimated covariance matrix, for large numbers of traits it can be necessary to reduce the actual number of parameters, however regularised they may be. A commonly used technique for dimension reduction in high-dimensional multivariate statistics is latent factor analysis (also called factor analytic), which is based on a rank-reduced representation of a covariance or correlation matrix. To explain the method, we first reparametrise the full MR-PMM (8) as

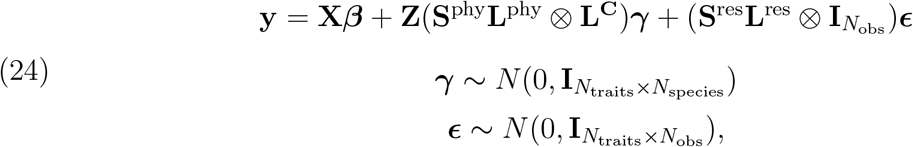

where **S**^phy^ and **S**^res^ are diagonal matrices of standard deviations and **L** are lower triangular Cholesky factors satisfying **R**^phy^ = **L**^phy^(**L**^phy^)^T^, **R**^res^ = **L**^res^(**L**^res^)^T^, and **C** = **L**^**C**^(**L**^**C**^)^T^, with **Σ**^phy^ = **S**^phy^**R**^phy^(**S**^phy^)^T^ and **Σ**^res^ = **S**^res^**R**^res^(**S**^res^)^T^; see Section 7.1 for definitions of **S** and **R**.

For the model (24), the rank of the correlation matrix **R**^phy^ (or **R**^res^) can be reduced from *N*_traits_ to *k* by replacing the *N*_traits_ *× N*_traits_ lower triangular square matrix **L**^phy^ with the rectangular *N*_traits_ *×k* lower triangular matrix **L**^phy(*k*)^ and ***γ*** with 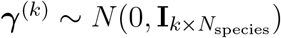, for a choice of integer *k < N*_traits_. The elements of **L**^phy(*k*)^ and ***γ***^(*k*)^ are called the *loadings* and *latent factors*, respectively.

It is important to note a few technical aspects of fitting latent variable models. Here we have chosen the *k*-column matrix of loadings to be a lower triangular matrix. It must also satisfy the constraints that the Euclidean row norms (i.e., the sum of squares of elements in each row) equals one and that the diagonal elements are positive valued. These constraints ensure that **L**^(*k*)^(**L**^(*k*)^)^T^ is a valid correlation matrix and that each matrix element is identifiable. These constraint choices are not unique and it is common to relax the lower-triangular restriction and require only that the columns be orthogonal. The latter allows all traits to contribute loadings to each factor which can aid direct interpretation of the factors. This choice, however, comes at the cost of sampling efficiency and the introduction of additional technical issues such as label switching (Hassler et al., 2022). A possible solution is to retain the lower triangular parameterisation for efficient model fitting and to compute the leading *k* eigenvectors of the estimated **L**^(*k*)^(**L**^(*k*)^)^T^ which are orthogonal and comprise contributions from all traits. As for the full rank case discussed in the previous section, regularising priors can be placed on factor loadings (Runcie and Mukherjee, 2013).

Latent factor models provide an effective means to reduce the complexity of high-dimensional multivariate models. In the MR-PMM setting, these models can be fit with the R package `asreml` (Butler et al., 2023) using residual maximum likelihood or using the Julia package `PhylogeneticFactorAnalysis`(Hassler et al., 2022) which acts as an interface for the Bayesian phylogenetic inference software `BEAST` (Suchard et al., 2018). Both packages offer parameterisation options and the latter will automatically select the number of factors *k* using cross validation. Neither `MCMCglmm` nor `brms`, the two packages on which we focus for tutorial, are able to fit latent factor models at this stage, although this class of models can be coded explicitly in the `stan` (Stan Development Team, 2024) statistical modelling language, which is the platform underpinning `brms`.

## 8 Discussion

### 8.1 Summary

Phylogenetic mixed models are a familiar, flexible, and scalable framework for comparative analyses of species traits. Phylogenies are used to derive an expectation of similarity among species trait values, which enters the model as a phylogenetic correlation matrix. Different evolutionary models are fit via transformations of this matrix, or else by deriving expected pairwise phylogenetic correlations under the chosen model. For a single response trait, modelling phylogenetic structure means the proportion of variance attributable to phylogeny (i.e., phylogenetic signal) can be estimated. However, single-response models with fixed effects may confound an ensemble of correlations if predictor and response variables both show phylogenetic signal. MR-PMM provides a solution via the explicit decomposition of trait covariances. This allows correlations between response traits to be partitioned into phylogenetic and non-phylogenetic contributions, providing a more detailed and informative breakdown of trait covariances. We review these models, with a focus on statistical concepts, model specification, parameter interpretation, and practical implementation. We discuss recent advances in methodology and propose a synthesis integrating multilevel distributional models with MR-PMM as a valuable direction for future work. Finally, we provide an extensive tutorial featuring worked examples and supplementary code to help users get started, as well as additional mathematical details for those seeking a more technical understanding.

### 8.2 Strengths and Weaknesses

One limitation of MR-PMM, as currently implemented in the popular R packages `MCMCglmm` and `brms`, is the restriction to a multivariate BM model of evolution. Despite its statistical elegance, BM is clearly a gross oversimplification of the varied and dynamic processes generating phenotypic diversity over macro-evolutionary time (Uyeda et al., 2018; Losos, 2011; Freckleton and Harvey, 2006). Most notably, BM does not explicitly account for adaptive evolution (e.g. stabilizing and directional selection). At first glance, this appears to be a fatal weakness. However, phylogenetic signal consistent with BM is, in fact, also consistent with adaptation via niche conservatism; these processes can produce indistinguishable phylogenetic patterns in ecological data (reviewed in Westoby et al. 2023; Wiens et al. 2010). Thus, even Brownian phylogenetic (co)variance is relevant for understanding divergence in ecological strategy between clades, a major contributor to the functional diversity of ecosystems.

Looking beyond PMM, there exists a broad class of processes expressible in terms of stochastic partial differential equations that characterise more complex and dynamic multivariate evolutionary models (Blomberg et al., 2020; Bartoszek et al., 2012; Butler and King, 2004). These include fully parameterised multivariate OU models (i.e., unstructured ***A*** matrices), as well as multiple optima and shift models (see below), examples of which are implemented in `mvSLOUCH` (Bartoszek et al., 2012), `mvMORPH` (Clavel et al. 2015) and `Rphylopars` (Goolsby et al., 2017), with some models also estimable via a fast linear-time likelihood algorithm in `PCMbase` (Mitov et al., 2020). Like MR-PMM, many of these methods accommodate missing data and within-species variation but offer more detailed and (potentially more) realistic evolutionary models. On the other hand, model selection approaches to identify the correct structure (e.g, for ***A***) are non-trivial, and may require very large comparative datasets (Bartoszek et al., 2023). Furthermore, these methods are restricted to continuous traits (though see Hassler et al. 2022) and do not permit an arbitrary multilevel decomposition of trait covariances, which is a desirable and informative simplification for many biological questions.

Another important limitation of the MR-PMM specified in (9), is that it assumes the chosen evolutionary model is both constant through time and homogeneous across the tree, making it vulnerable to miss-specification under certain evolutionary scenarios, e.g., rare events (Uyeda et al., 2018; Clavel et al., 2015). Some alternative methods allow for shifts, meaning parameters of the evolutionary model (e.g. 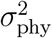), even the model itself (BM, OU, etc), are allowed to vary between clades and across time (Pagel et al., 2022; Mitov et al., 2020, 2019; Clavel et al., 2015), including (when furnished with fossil data) changes in the trait-level covariance matrix itself through time (i.e., Σ = Σ(*θ, t*)) (Blomberg et al., 2020). Currently, it is not clear how these more dynamic models of trait evolution might be expressed in the PMM framework. Thus, other techniques will be more appropriate when trying to infer the exact timing or phylogenetic position of rare evolutionary events or shifts in the underlying processes generating trait variation (Pagel et al. 2022; Uyeda and Harmon 2014; Slater 2013).

One strength of the mixed model approach is their extension to generalised linear models that permit multivariate (Gaussian) random effects on the link-transformed mean of traits that are themselves not Gaussian distributed (Nakagawa and Schielzeth, 2010; Hadfield, 2010). Many evolutionary hypotheses invoke causal relationships between continuous and discrete traits. However, currently MR-PMM is one of few methods available to estimate correlations between continuous and discrete traits in a fully phylogenetic framework (see also Haba and Kutsukake 2019). Despite complications arising from non-linear latent variable transformations (see Section 3.0.5), this has enabled several long-standing evolutionary hypotheses to be addressed in recent years (e.g., Downing et al. 2020; Cornwallis et al. 2017). A second strength of MR-PMM is the capacity to specify multi-level models that appropriately capture correlated hierarchical effects (Cinar et al., 2022; Nakagawa and Santos, 2012; Hadfield and Nakagawa, 2010). Integrating data from multiple studies to draw general conclusions is a common practice in comparative biology. It enables researchers to increase the power and complexity of analyses and is simply necessary to make full use of growing public data repositories. The capacity to account for multi-level structure is therefore crucial for methods to effectively scale with modern data compilations (e.g. Falster et al. 2021; Kattge et al. 2020).

### 8.3 Recommendations and Future directions

One challenge many analysts face when attempting a phylogenetic comparative analysis is that not all available species with trait data are featured in a suitable published phylogeny. A powerful workaround is to use the best available species-level phylogeny to derive a phylogeny for a higher taxonomic rank (e.g., series, section, or genus). This approach has two tangible benefits: 1) computations involving the phylogenetic correlation matrix **C** are a major bottleneck in the evaluation of the model likelihood. Therefore, reducing the dimension of **C** by refocusing to a higher taxonomic rank comes with substantial reductions in model fitting time; 2) It is often possible to include data from more species, as those missing from species-level phylogenies can instead be treated as replicates of a higher taxonomic rank that does feature in the phylogeny. This method has the obvious drawback of ignoring phylogenetic structure close to the tips (e.g., sister species relationships), modelling only those effects owing to deeper splits within the tree. However, in cases such as the Eucalypts, where taxonomic series often represent freely hybridising species groups (Pfeilsticker et al., 2023; Larcombe et al., 2015), modelling phylogenetic effects above the species level may in fact be desirable, as it relaxes the assumption that recent divergences can be expressed in terms of a strictly bifurcating evolutionary process.

A more general pitfall in phylogenetic comparative analyses is the failure to adequately account for multilevel structure. Assuming sufficient replication, the preferred approach is always to model species traits at the lowest possible level (e.g., observations on individual organisms), using a multilevel model. Partitioning variation across the model hierarchy utilises all available information, while facilitating evaluation of more interesting biological questions. For example, separation of within- and between-group variance components (e.g., species, population, individual) (Garamszegi and Møller, 2017; Westneat et al., 2015); separation of phylogenetic and spatial effects (Gomes et al., 2023; Freckleton and Jetz, 2009); and the estimation of within-species residual (co)variances (i.e., patterns of species-specific phenotypic plasticity and trait co-variation) (Goolsby et al., 2017). Detailed variance partitioning may, in fact, be necessary to identify drivers of selection in natural populations. For example, partitioning variance in anti-predator behaviour of 254 bird species supported the view that within- and between-population variances are driven by different selective forces (Garamszegi and Møller, 2017). Similarly, within-species variances account for a considerable proportion of community-level variation in plant functional traits, particularly for whole-plant (e.g. plant height), rather than organ-level (e.g. leaf size), traits (Siefert et al., 2015). As species differ not only in mean trait values but also in the magnitude and structure of trait variation, these studies highlight the enormous potential of multilevel distributional models to advance our understanding of the processes producing and maintaining biological variation.

Integrating multilevel distributional MR-PMM with sophisticated evolutionary models remains a considerable methodological challenge (see section 3.0.4). In particular, software implementations are limited and modern comparative analyses including many thousands of taxa pose genuine computational barriers to more complex, parameter-rich evolutionary models. Of course, such challenges do not obviate the need to account for structural dependencies in comparative data, rather they call for a more tractable model. Of the available options, BM is easily the most tractable, a virtue so often repeated it risks being overlooked (Blomberg et al., 2020). Nonetheless, large-scale studies, particularly in the field of plant functional ecology, continue to be published without directly accounting for phylogeny (e.g., Joswig et al. 2022; Bruelheide et al. 2018; Díaz et al. 2016; also see Koricheva and Gurevitch 2014). At the very least, phylogenetic models assuming BM evolution should feature in the analysis of such datasets, and MR-PMM offers a robust, flexible and informative framework to do so (Westoby et al., 2023). However, with the inexorable expansion of biological trait databases (Falster et al., 2021; Kattge et al., 2020), even models assuming BM can become computationally burdensome as the number of species and traits included in analyses grows. Future work could address this by developing approximate inference methods for MR-PMM, such as mean field variational Bayes algorithms (Ormerod and Wand 2010, though the accuracy of covariance components for non-Gaussian response traits remains a challenge, see Hughes et al. 2023). A particularly powerful approach would emerge from integrating recent advances in the computational efficiency of non-Brownian multivariate evolutionary models (Mitov et al., 2020) into the generalizable, multi-level framework of MR-PMM.

Another clear direction for future research is a more comprehensive assessment of statistical power and sensitivity in MR-PMM. In particular, it would be instructive to have clearer expectations around parameter uncertainty with respect to the number of response traits, the mixture of response distributions, the relative magnitude of phylogenetic and non-phylogenetic (co)variances, as well as phylogenetic uncertainty and sample sizes. In general, MR-PMM is well suited to datasets with many species *n* relative to the number of traits *p* (i.e., high *n* : *p* ratios). Reduction of model complexity, for example via shrinkage (Section 7.1) or rank-reduction of the covariance matrix (Section 7.2), may be required for data sets with low *n* : *p* ratios (e.g., geometric morphometrics), for which specialised methods are also available (Clavel and Morlon 2020; Adams and Collyer 2019, 2018; Collyer et al. 2015; Adams 2014; Hassler et al. 2022, also see Runcie et al. 2021 for methods relevant to genomic prediction). However, a formal examination of power and sensitivity, provided by simulation studies, is necessary to clarify the inherent limitations of estimation and inference in an MR-PMM framework. Fortunately, the ability to impute missing response values often means that more data can be included in analyses. Even relatively small data sets can potentially yield useful inferences by using sparsity-inducing priors (Section 7.1) to identify a small subset of parameters that escape shrinkage to zero.

### 8.4 Conclusions

1. Species phenotypes are the product of a complex network of causal processes that promote co-selection of traits, over both short and long time scales. For some traits, (co)variation is conserved over evolutionary time, creating patterns in inter-species data that reflect phylogenetic relationships (Section 2).
2. Distinguishing the influence of these conserved effects from those that are decoupled from phylogenetic history is a fundamental objective of evolutionary ecology, because it is the balance of these forces that defines whether constraints, trade-offs and coordinated strategies, should be understood in terms of deep evolutionary integration or labile responses to prevailing conditions (Section 4).
3. Univariate approaches to cross-species analysis are not parameterised to distinguish between these components of trait correlation (Section 2), leading to potential confounds and inferential limitations. In contrast, MR-PMM, in which correlations between traits can be partitioned into phylogenetic and non-phylogenetic components, offer a more plausible and informative model of trait evolution under many circumstances (Section 4, 6).
4. The capacity to fit multilevel distributional models (3.0.1, 3.0.3), continuous and discrete response variables (3.0.5), estimate partial correlations (3.0.7), impose different models of trait evolution (3.0.4), and employ conditional predictions for imputation and model validation (5), makes MR-PMM a robust and unifying approach to many open questions in comparative biology.
5. In particular, MR-PMM is well-equipped to advance an emerging synthesis between genetic, ecological, and evolutionary approaches to the study of trait variation. Data sets will soon exist that allow the partitioning of within- and between-species trait variances into additive genetic, phylogenetic, spatial, environmental, and residual components. We argue that careful parameterisation of MR-PMM (Sections 7.1, 7.2) may offer a scalable framework for analysing these large complex datasets.
6. Currently, software implementations of the various methods discussed are available piecemeal across several different applications. In practice, we found this led to different use cases for the two R packages we explored for fitting MR-PMM. The analytic solutions leveraged by a Gibbs sampling approach, give`MCMCglmm` a considerable computational advantage when all responses variables are Gaussian with the restriction that conjugate priors be used. In contrast, the Hamiltonian Monte Carlo sampler used by `brms` is likely to outperform for non-Gaussian error distributions and allows flexible prior specification. `brms` also allows for distributional models, benefits from a user-friendly syntax and integrates well with other useful packages such as `tidyverse` and `loo`.
7. We expect more researchers in ecology and evolution will adopt MR-PMM as barriers to implementation for non-specialists are broken down. To this end, we provide extensive tutorial materials, including annotated code and applied examples.

## 9 Tutorial

Tutorial material relevant to this article can be found here.

## 10 Acknowledgements

We thank Ian Wright, Mark Westoby, David Warton, and Simone Blomberg for helpful discussions during preparation of the manuscript. We thank Shinichi Nakagawa for editorial suggestions, and Jarrod Hadfield and Pierre de Villemereuil for detailed reviews that greatly improved the manuscript. We thank Rachael Gallagher for consultation on the AusTraits database. This work was funded by The Australian Research Council Centre of Excellence for Plant Success in Nature and Agriculture (CE200100015).

